# Scale-dependent associations between genetic similarity and aggregation behaviour in three toad lineages

**DOI:** 10.1101/453316

**Authors:** Kazuko Hase, Masato S. Abe

**Affiliations:** Department of General Systems Studies, The University of Tokyo, Meguro, Tokyo 153-8902, Japan; Graduate School of Global Environmental Studies, Kyoto University, Yoshidahonmachi, Sakyo-ku, Kyoto, 606-8501, Japan; Graduate School of Culture and Information Science, Doshisha University, Kyotanabe, Kyoto, 610-0394, Japan; Center for Advanced Intelligence Project, RIKEN, Chuo-ku, Tokyo, 103-0027, Japan; CBS-TOYOTA Collaboration Center, RIKEN, Wako, Saitama, 351-0198, Japan

**Keywords:** aggregation behaviour, genetic relatedness, kin recognition, major histocompatibility complex (MHC) gene, toad tadpole

## Abstract

Social aggregation is widespread in vertebrate larvae, yet how genetic relatedness shapes group cohesion remains poorly understood. In particular, the association between genetic similarity and social aggregation may depend on the scale at which genetic relatedness is evaluated. Here, we quantitatively examined aggregation behaviour in tadpoles from three lineages of Japanese toads, *Bufo japonicus*, *B. formosus*, and *B. gargarizans miyakonis*. We experimentally combined tadpole cohorts differing in relatedness and lineage and quantified aggregation using spatial dispersion analyses. We then examined relationships between aggregation behaviour and genetic variation using mitochondrial DNA (mtDNA), major histocompatibility complex (MHC) class II variation, and microsatellite diversity. Overall degree of aggregation did not differ significantly between sibling and non-sibling cohorts, and Morisita’s index provided no evidence that tadpoles formed separate clumps according to relatedness. However, the correlation between genetic variation and aggregation differed across scales of genetic relatedness: among non-sibling cohorts, greater mtDNA pairwise distance was associated with weaker aggregation, whereas among sibling cohorts, greater MHC amino-acid difference was associated with weaker aggregation. These results indicate that the degree of aggregation is related to genetic similarity across different scales of relatedness, rather than being explained by a simple sibling–non-sibling distinction. Our findings suggest that tadpole group cohesion is related to genetic similarity in a scale-dependent manner, highlighting the importance of considering the scale of genetic relatedness when investigating social discrimination in vertebrates, while the sensory and molecular mechanisms underlying these relationships remain to be determined.

## INTRODUCTION

The evolution of cooperation is a major issue in evolutionary biology. Since the proposal of kin selection theory [1,2], social behaviours have often been discussed in relation to differences in genetic relatedness among interacting individuals. Kin recognition can influence social interactions in some animals, although kin discrimination is not required for an action to have indirect consequences on fitness. As limited dispersal ability leads to high relatedness among interacting individuals, kin selection theory could explain cooperation to some extent even in the absence of kin recognition ability [2]. In viscous populations, due to the population structure behaviours that benefit neighbours can have beneficial effects on relatives. Under certain population conditions, “help by neighbour” cooperation can evolve without any information on relatedness; however, kin recognition systems have been discovered many times across taxa, from unicellular organisms to humans [3, 4].

Despite exhibiting low dispersal ability, anuran amphibians have been tested for kin recognition abilities in terms of both larval behaviour [5–10] and mating preferences [11].

Tadpoles commonly aggregate into groups and, similar to fish, their schooling behaviour varies between centralised for predator avoidance and non-centralised for foraging [12–16]. Cohesive aggregation of toad tadpoles has been proposed to reduce predation risk and increase survival [14,16]. Groups of the tadpoles are usually derived from multiple clutches, such that some individuals are siblings and others are not [17]. Although the tadpole groups can therefore contain individuals with different degree of genetic relatedness, it remains unclear exactly how genetic structure affects aggregation behaviour [16]. In other words, we do not understand how genetic background relates to social aggregation.

Previous studies of kin recognition have often examined whether individuals preferentially associate with kin rather than non-kin, implicitly linking social discrimination to group membership or group favoritism [18]. The degree of social discrimination may instead vary continuously with genetic similarity, as shown in social amoebae [19]. The influence of genetic relatedness on social aggregation may vary across species and across different ranges of genetic relatedness. In some species of true toad (*Bufo americanus*, *B. boreas*, *B. bufo*, *B. melanostictus*, and *B. scaber*), tadpoles tend to aggregate more cohesively in siblings more than non-siblings [5,6,14], suggesting that genetic relatedness may influence social association in some amphibians [20,21]. In contrast, *Bufo formosus* did not exhibit kin-biased association in a recent binary-choice test, regardless of early social experience or water environment [22].

Genetic similarity could instead be related to the cohesion of mixed groups, even when individuals do not form distinct kin-specific groups. This possibility raises the question of whether genetic similarity is related to group cohesion across different scales of genetic relatedness.

*Bufo japonicus* (hereafter, *japonicus*), *B. formosus* (*formosus*), and *B. gargarizans miyakonis* (*miyakonis*) are native Japanese toads belonging to the true toad family, Bufonidae. Whilst the distribution of these toads is isolated geographically across the islands of Japan (*japonicus* and *formosus* are distributed in the western and eastern regions of the mainland and *miyakonis* inhabits the Miyako Islands), the ecological features of these toads are somewhat common [23]. Genetic differentiation among these Japanese toads is related to their geographic distribution [24,25]. According to classification based on mitochondrial DNA (mtDNA) sequences, they belong to three distinct clades [26]. In contrast to the similarities between *japonicus* and *formosus*, adult *miyakonis* shows clear differences in morphology, possessing a smaller size and fewer bumps [23]. However, the larvae of these toads are morphologically similar. All tadpoles are basically herbivores and exhibit aggregation behaviour.

Here, we asked whether the relationship between genetic similarity and tadpole aggregation depends on the scale of genetic relatedness. We tested aggregation among sibling and non-sibling cohorts representing different degrees of genetic relatedness, including pairs within and between mtDNA lineages. We quantified genetic variation using mtDNA sequence divergence, MHC class II amino-acid differences, and microsatellite diversity, and examined whether these measures were associated with the strength of aggregation. MHC genes not only play a central role in the immune system, but have also been implicated in kin and social recognition in several vertebrates [27–34]. MHC-associated variation has been proposed to contribute to individual odour cues involved in social recognition [35,36]. We predicted that, if genetic similarity contributes to social aggregation, it should increase as genetic diversity decreases. We further asked whether the genetic predictors of cohesiveness differ between more closely and more distantly related cohorts, indicating scale-dependent associations between genetic relatedness and social aggregation.

## MATERIAL AND METHODS

### Experimental animals

Between February and March 2013, we sampled 18 clutches from eight reproductively isolated sites in Japan (for details, see electronic supplementary material Table S1 and Figure S1). The clutches belonged to three different lineages of Japanese toad: *japonicus* (*n* = 7), *formosus* (*n* = 8), and *miyakonis* (*n* = 3). Clutches spawned by amplectant pairs consisting of a single male, as far as we observed until spawning finished, were specifically chosen to avoid polyandry. After the amplectant pair had left the spawned egg strings, we collected approximately 0.4 m of egg strings containing about 200–400 embryos and transported them to the laboratory. According to a polyandry assessment based on microsatellite markers, 1 of the 18 clutches (NK) showed the possibility of two sired males (see below and the supplemental Experimental Procedures Method [EPM]). Hence, in this paper, sib does not mean full-sibling, and the sib cohorts of the NK clutch were actually half-siblings.

Embryos belonging to the same clutch were divided into two sets before hatching (100–150 individuals per set) and transferred into a container (220 mm width × 310 mm diameter × 40 mm height) in a large incubator at 18°C under a 12:12 h dark:light cycle. All tadpoles were raised with respect to each set with a sufficient supply of fish food pellets containing vegetable stock as the main component (PLECO; Kyorin, Hyogo, Japan).

Individuals of each set were colour-marked by immersion in either a 0.00025% solution of neutral red for 24 h or a 0.00025% solution of methylene blue for 12–24 h before the aggregation test, as described previously [37]. Thus, sib tadpoles from the same clutch were randomly distributed into different colour sets (red or blue) without harm or the addition of exogenous odour. This treatment was used simply to identify clutch type in the duplication of the aggregation test and genotyping. After the aggregation test and genotyping (see below), the tested tadpoles were released into their native ponds when possible, excluding *miyakonis*, which is an artificially introduced alien species.

The total DNA content was extracted from the tip of the tails of all tadpoles included in the aggregation tests. The mtDNA of all clutches was analysed using DNA sequencing, as described previously [37]. MHC class II antigen (partial exon) allele sequences were amplified by PCR using previously developed locus-specific primers [39]. Then we genotyped all of the tadpoles included in the test. Thus, a total of 3,400 individuals were genotyped using restriction fragment length polymorphism analysis of PCR-amplified fragments. To estimate the genetic diversity in the sib group, individuals were genotyped using four microsatellite loci that showed polymorphisms across 18 clutches: *Bbufu23*, *Bbufu39*, *Bbufu62* [40], and *Bjap14* [41]. The DNA extraction and genotyping methods are described in the EPM and Tables S1 and S2.

### Aggregation behaviour test

An aggregation behaviour test was conducted in 68 sets consisting of 50 randomly chosen tadpoles from the different laboratory-reared sets (25 red- and 25 blue-dyed tadpoles). The tadpoles were transferred into plastic containers (230 mm width × 350 mm diameter) containing 1 L dechlorinated tap water (20 mm in depth). Pairs of red- and blue-dyed tadpoles were characterised as sib (i.e., derived from the same hatch but raised separately) or non-sib (i.e., intra-/inter-lineages). The aggregation behaviour test consisted of three trials using the same cohort of 50 tadpoles. The selection process lasted for 5 days. The aggregation tests were performed according to the procedure described below (also shown in Figure 1). Tadpoles were chosen for a trial cohort according to their developmental stage to ensure that each cohort consisted of tadpoles of a similar age belonging to stages 27–37 [42]. The trials proceeded with reference to the methods of a previous study [5]. All trials lasted for 24 h in a closed large incubator. Each trial started at night. After 50 tadpoles were individually evenly distributed throughout the container using a spoon, as pre-conditioning, the container was kept in the dark for 12 h. We began observing the aggregation behaviour the following morning. Aggregation levels were recorded under normal light conditions using a Pentax Optio W80 camera (Ricoh, Tokyo, Japan) in interval shooting mode. Photographs were taken at 1 min intervals, yielding a total of 720 shots per trial. During the trials, the containers with the tadpoles were maintained in large incubators at 18°C for 24 h. During this period, the tadpoles were not fed or interrupted (i.e., the incubator door remained closed for the duration of the test). After completing this first trial, the tested cohort was divided into two sets based on the colour of the dye (25 red tadpoles and 25 blue tadpoles) and placed on small plastic containers half the size of the test container (230 mm width × 155 mm diameter × 40 mm height), fed, and allowed to acclimate for 24 h (12:12 h dark:light) at 18°C (intermission process). Then the tadpoles were returned to the experimental container for a repeated aggregation trial, as described above. The same procedure (intermission and trial) was repeated once more. Overall, the experimental process included three tests and two intermissions (trial, intermission, trial, intermission, trial). Figure 1 presents an outline of the experimental process. Aggregation tests were conducted in 68 cohorts consisting of two laboratory-reared set pairs, including 26 sib pair cohorts (*japonicus*, *n* = 10; *formosus*, *n* = 10]; *miyakonis*, *n* = 6), 17 non-sib intra-lineage cohorts (*japonicus* + *japonicus*, *n* = 5; *formosus* + *formosus*, *n* = 9; *miyakonis* + *miyakonis*, *n* = 3), and 25 non-sib inter-lineages *japonicus* + *formosus*, *n* = 11; *japonicus* + *miyakonis*, *n* = 8; *formosus* + *miyakonis*, *n* = 6).

**Figure 1.**
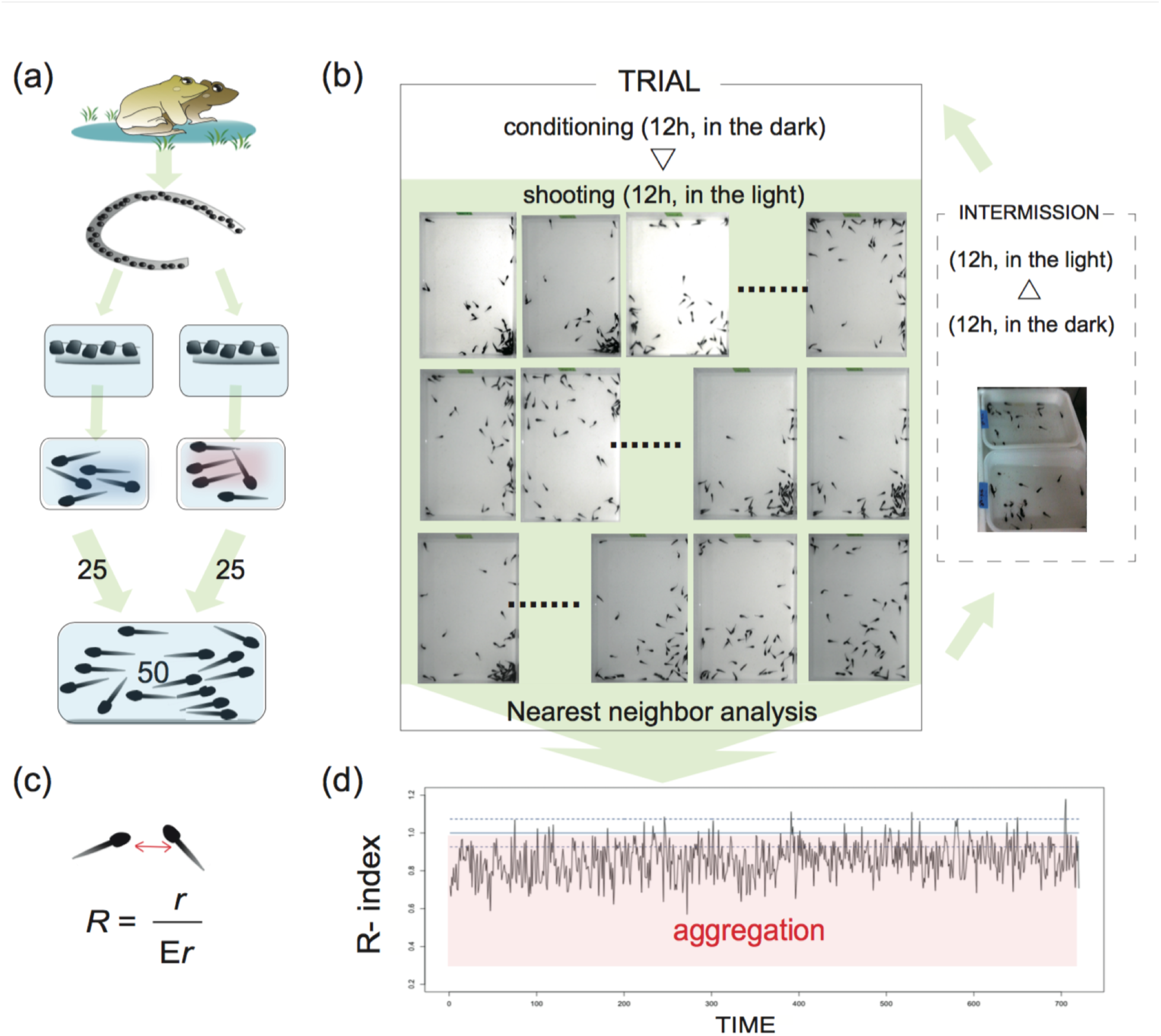
(**a**) Diagrammatic illustration of the experiments: from sampling to aggregation test. (**b**) Practical example of the test. (**c**) Outline of the nearest-neighbour analysis. (**d**) Example of analysis (among 68 × 3 trials).

### Aggregation index

The degree of aggregation was measured as the aggregation index (R index) based on the nearest-neighbour analysis [43, 44]. We obtained the positions of 50 individuals in the two-dimensional (2D) arena (230 mm × 350 mm), after which we calculated the mean distance *r* from each individual to its nearest neighbour. For randomly distributed individuals, the mean distance (*E_r_*) from the nearest individual was analytically described [43]. However, to incorporate the effects of the corners of the area, we conducted stochastic simulations, scattering 50 random points within the arena. After 1,000 iterations, we obtained *E_r_* ± standard deviation (SD) = 21.11 ± 1.7 (mm), with a lower 5% confidence limit of 18.28 (mm). When *r* was below 18.28 (mm), the distribution of the tadpoles could be ascribed to statistically significant aggregation behaviour (R < 1). It should be noted that lower R index values indicate higher degrees of aggregation. In addition, we defined the time as the aggregation time with a lower confidence limit (5%) over three trials, presented as the number of images per 720 images. To assess whether there were differences in aggregation formation (e.g., number of clumps) between sib and non-sib cohorts, we measured the distribution patterns in the image showing the greatest degree of aggregation for each tested cohort using the Morisita index (*I_δ_*) [45]. To apply this index, we divided the 2D arena into 16 sections and counted the number of tadpoles in each section (Figure S2). High *I_δ_* values indicated greater spatial clumping of tadpole. After the analyses, we compared the scores calculated from the R index and *I_δ_*. Thereafter, we used the R index as the aggregation index.

### Tadpole gene diversity

We used three indices to compare the gene diversity among the 68 cohorts: mtDNA pairwise distance, MHC (class II antigen) amino acid sequence difference, and microsatellite diversity (only for sibs) based on the four aforementioned loci. Although mtDNA and microsatellites were used as neutral genetic markers, MHC was used as an adaptive genetic marker potentially relevant to kin recognition. After analysing the phylogenetic relationships (electronic supplementary material EPM1 and Figure S3), we quantified the genetic diversity of each marker.

The evolutionary divergence of the mtDNA lineages for each experimental clutch pair was estimated using the number of base substitutions per site. These analyses were conducted based on the maximum composite likelihood model [46] in MEGA 5.2 [47]. All values were used to estimate the genetic differences among the mtDNA lineages as a genetic index. In these three toad lineages, the differences in the mtDNA cytochrome b sequences reflected their geographic distribution in Japan [24,36]. In addition, because there was no difference in the haplotypes of the microsatellite loci between *japonicus* and *miyakonis*, we determined mtDNA sequences to be more suitable as a genetic index for the interpretation of the phylogenetic relationship across the 18 clutches. The MHC genotypes were derived from the MHC class II antigen (partial exon, 0.3 kb) haplotypes for each cohort based on 13 alleles that were identified by genotyping (electronic supplementary material EPM2 and Tables S1 and S2). We calculated the amino acid sequence differences of the MHC genotypes in each cohort as an additional genetic index [48] using MEGA 5.2 [47]. In addition, we adopted four loci of the microsatellite markers to assess the gene diversity in the sib cohort (electronic supplementary material EPM3). After genotyping, the microsatellite diversity score (average gene diversity over loci within the cohort) was calculated based on the number of alleles using Arlequin ver. 3.5.2 [49].

### Data analysis

To evaluate aggregation behaviour, the data were analysed with two-step analyses in R ver. 3.2.2 [50]. First, we used the two-way repeated measures analysis of variance (ANOVA) to assess the effects of relatedness (sib and non-sib [intra- or inter-lineages]) and the differences in lineages (affiliation of the mtDNA lineage of the tested pairs, e.g., *j*+ *j*, see Table 1). In this analysis, the mixed models approach was adopted using the lme4 package in R ver. 3.2.2 [51]. We treated relatedness as a fixed variable and mtDNA lineage as a random variable. In addition, it was conducted using values not only R index but also *I_δ_* as complement assessment. Second, to assess the relationship between genetic variation and aggregation behaviour, we used generalised linear models (GLMs; Gaussian distribution, identity function link). In the GLMs, we treated the R value as the response variable. In particular, GLMs were conducted to evaluate whether the three genetic indices were associated with the degree of aggregation. Considering the difference in the components of the genetic diversity among the tested cohorts, this analysis was performed at two levels: non-sib and sib cohorts. To identify elements exerting a significant effect on aggregation behaviour, we selected a candidate model based on the Akaike information criterion (AIC) differences (ΔAIC) and the Akaike weight (*w_i_*), with the weight of evidence in favour of model *i* as the best model in the set [51]. Explanatory variables were provided based on the tested cohort data (described in Table 1). The parameters for the comparison of explanatory models included MHC (amino acid sequence differences), developmental stage, mtDNA lineage affiliation of the tested pair (category: *j* + *j*, *f* + *f*, *m* + *m*, *j* + *f*, *j* + *m*, and *f* + *m*), mtDNA pairwise distance (for non-sib cohorts), the interaction effect of mtDNA and MHC (for non-sib cohorts), microsatellite diversity (for sib cohorts), and the interaction effect of microsatellite diversity and MHC (for sib cohorts).

**Table 1.**
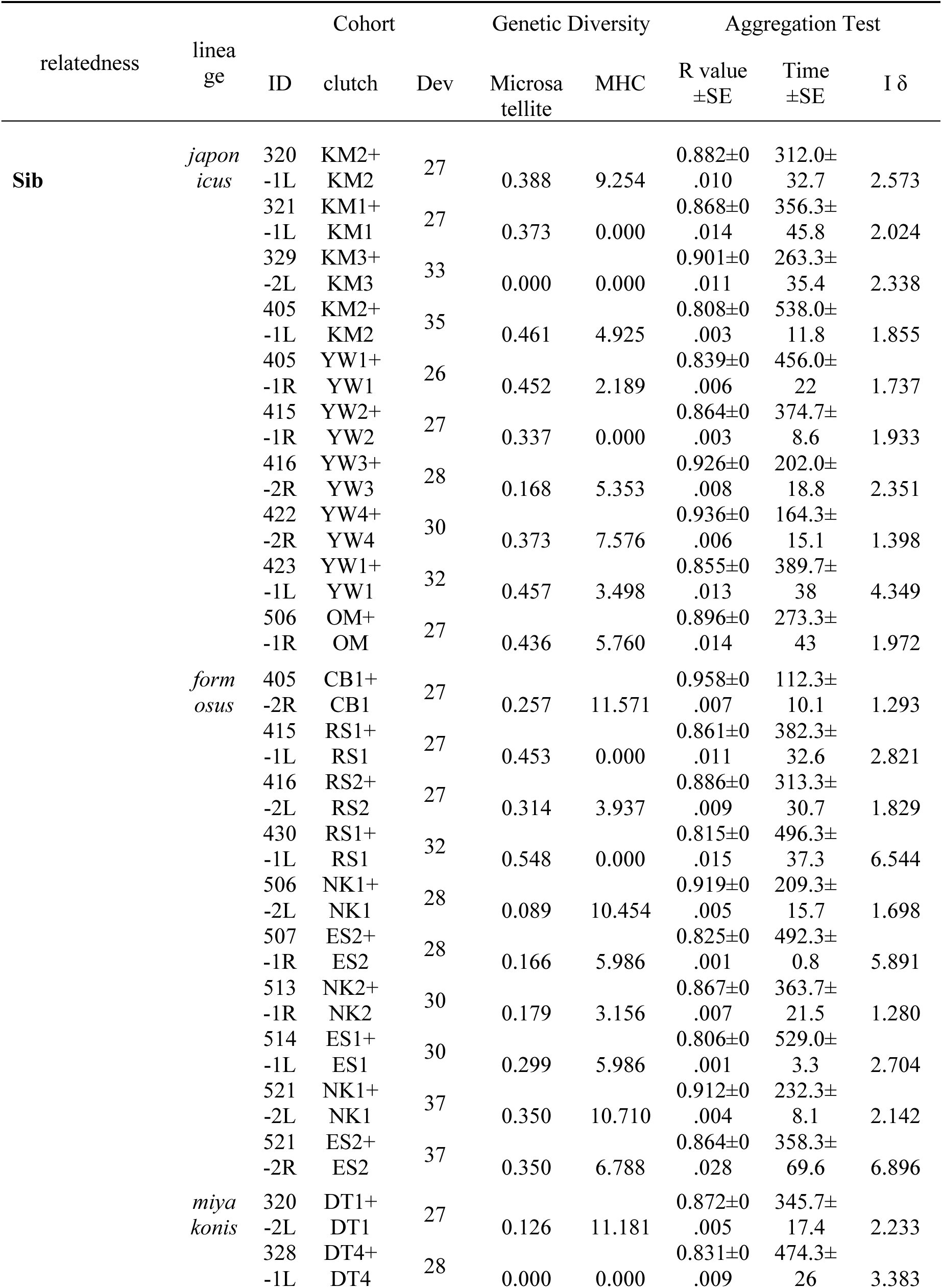
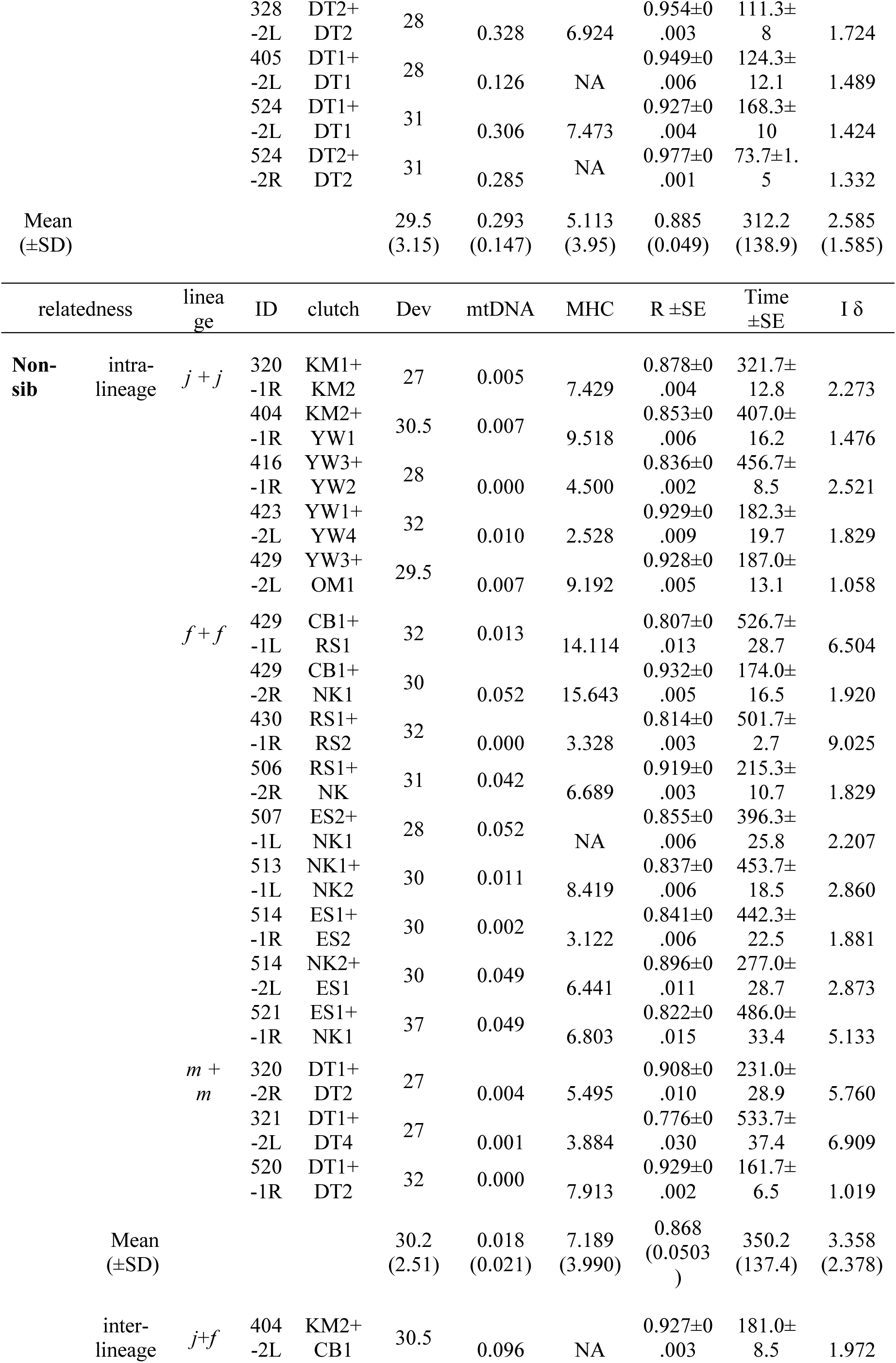
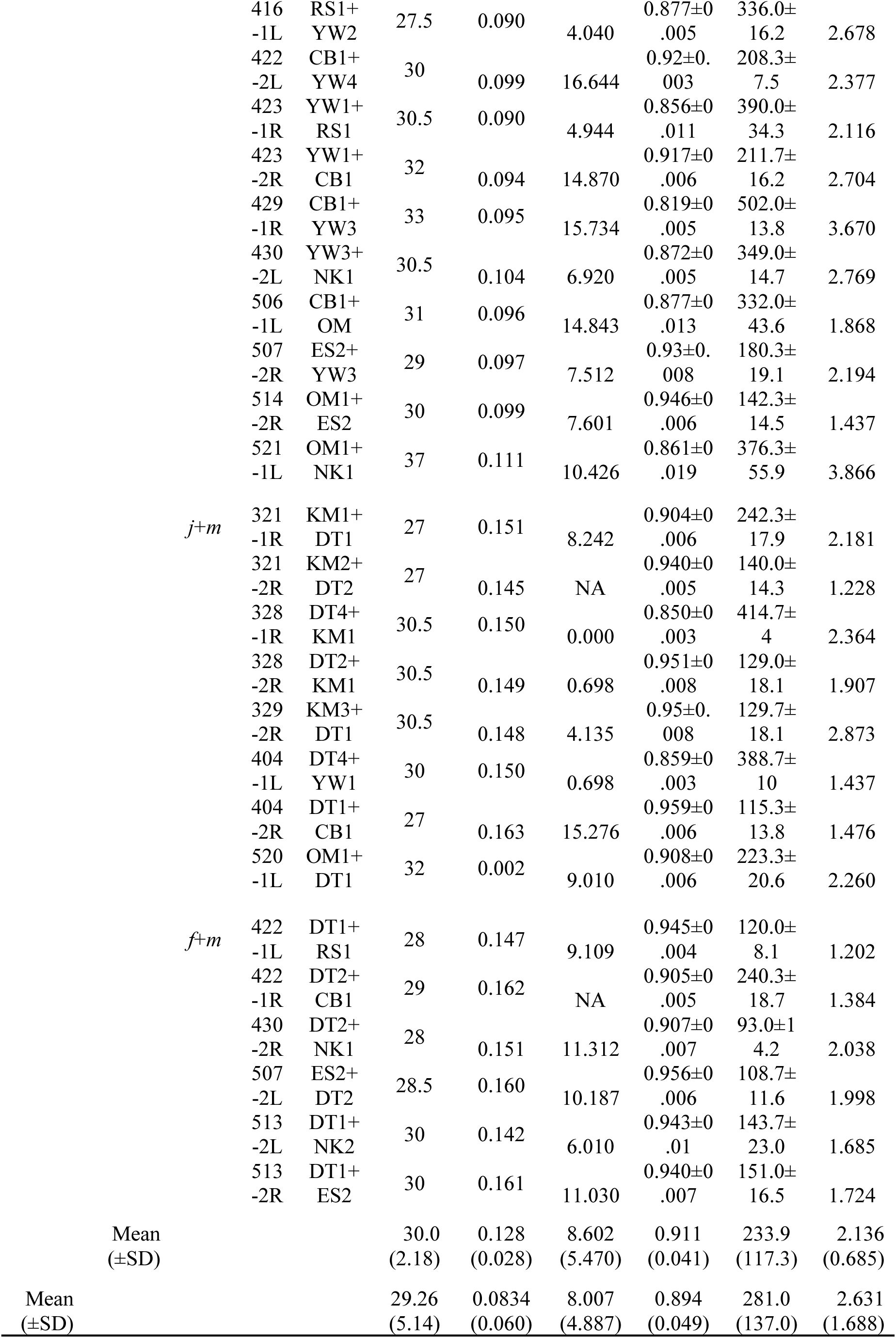
Results of the aggregation tests. The trial cohort information provided is as follows: relatedness, cohort ID, clutch affiliation for tested sets (detailed information is presented in Table S1 and Figure S1), lineage of tested clutch pairs (j: japonicus; f: formosus; m: miyakonis) and average developmental stages of tested tadpoles. Genetic diversity and aggregation test scores: mtDNA, pairwise distances; microsatellite, microsatellite diversity; MHC, amino acid differences by cohort. R value, average value of R index over 720 shots over three trials. Time, average aggregation time with lower confidence limit of 5%. Iδ, score of Morisita index [40].

## RESULTS

### Aggregation across lineage boundaries

We confirmed that tadpoles could aggregate across distinct lineage boundaries. We conducted aggregation behaviour tests on 68 cohorts, each consisting of 50 tadpoles that were raised separately in two sets (Figure 1). Table 1 shows the results of the aggregation test. In all tests, tadpoles showed significant aggregation behaviour: the R index, over 720 shots and three trials (hereinafter called the R value) was 0.8899 ± 0.0486 (*n* = 68, mean ± SD; range: 0.775–0.970); the aggregation times with lower confidence limits (5%) over three trials (hereinafter called Time) were 293.94 ± 136.24 min (*n* = 68, range: 93.0–538.0 of 720 min). In addition, there was a strong correlation between R values and Time (d.f. = 66, *P* < 0.0001, Pearson’s *r* = −0.995). Therefore, we predominantly used the R value to assess aggregation behaviour.

### Kin and lineage discriminations based on genetic relatedness

Our results showed no significant differences in the degree of aggregation among relatedness categories even considering differences in the lineages of the sib, inter-lineage, and intra-lineage cohorts (two-way repeated measures ANOVA: *F* = 2.356, d.f. = 2, *P* = 0.139, Figure 2). We did not detect any significant difference in R values between sib and non-sib (inter- or intra-lineage) cohorts, nor were there any significant differences in aggregation formation among the sib, intra-lineage, and inter-lineage cohorts based on *I_δ_* scores (two-way repeated measures ANOVA: *F* = 1.9616, d.f. = 2, *P* = 0.1693). The distribution patterns of the tadpoles did not differ depending on whether cohorts consisted only of sibs or admixed non-sibs (red- or blue-stained individuals). Moreover, *I_δ_* was correlated with the R value (d.f. = 66, *P* < 0.0001, Pearson’s *r* = –0.591, Figure S4).

**Figure 2.**
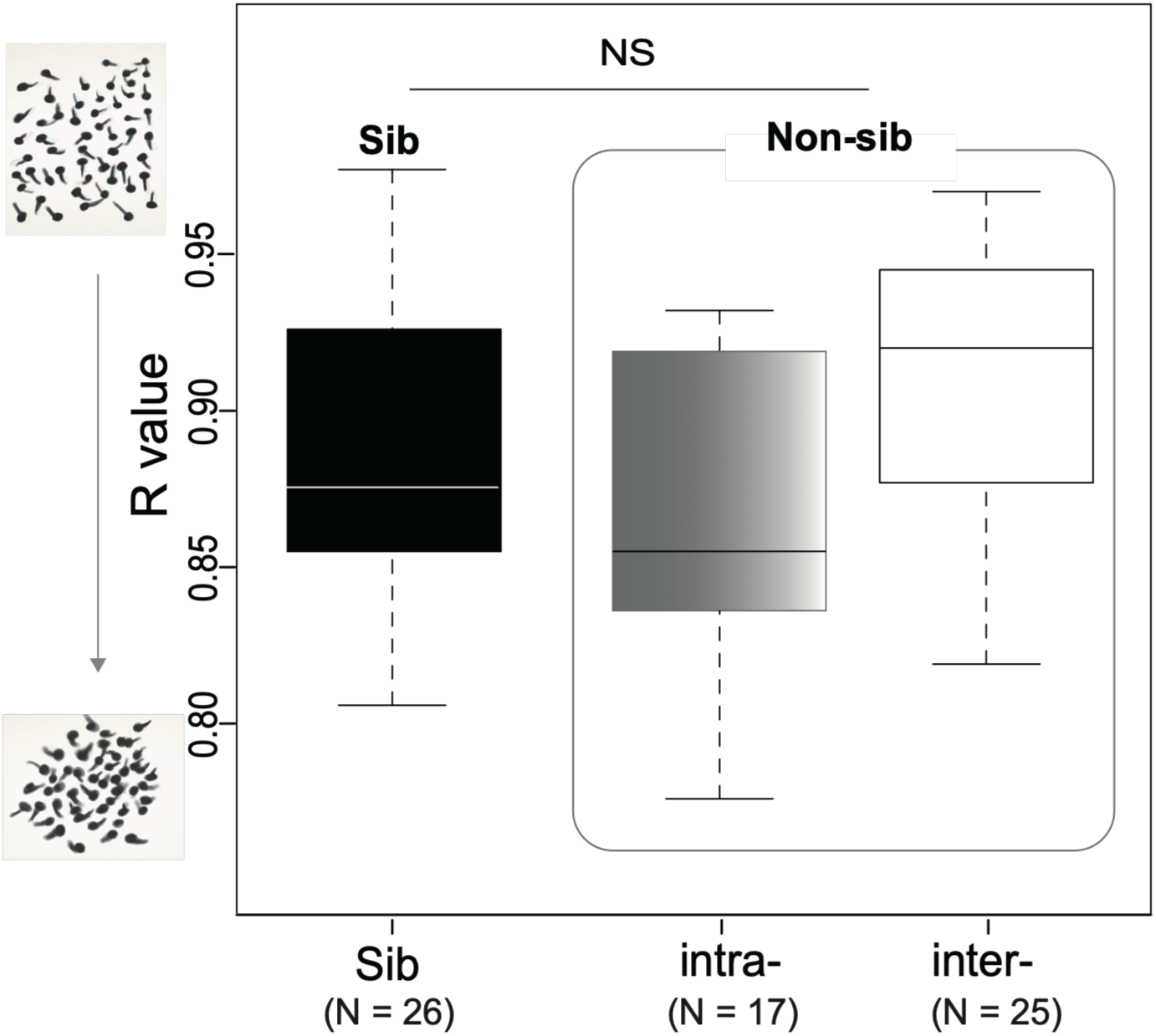
Comparison of aggregation values (R value) for relatedness between sib, inter- and intra-lineage pairs. Data by box plot (*n*, median, mean ± SD): sib cohort: 26, 0.877, 0.885 ± 0.049; intra-lineage cohort: 17, 0.872, 0.878 ± 0.050; inter-lineage cohort: 25, 0.920, 0.911 ± 0.041. There were no significant differences between the sib and non-sib (intra- or inter-lineage) cohorts (two-way repeated measures ANOVA: *F* = 2.356, d.f. = 2, *P* = 0.139).

Meanwhile, genetic variation was significantly associated with tadpole aggregation behaviour. In the non-sib cohorts, while mtDNA was significantly associated with R value (*n* = 42, Spearman’s *r* = 0.534, *P* < 0.001; Figure 3a), MHC amino acid differences were not associated with R value (*n* = 38, Spearman’s *r* = 0.231, *P* = 0.163; Figure 3b). In the sib cohorts, although microsatellite diversity was not correlated to R value (*n* = 26, Spearman’s *r* = −0.350, *P* = 0.079; Figure 3c), a relationship between MHC amino acid differences and R value was observed (*n* = 24, Spearman’s *r* = 0.573, *P* = 0.003; Figure 3d).

**Figure 3.**
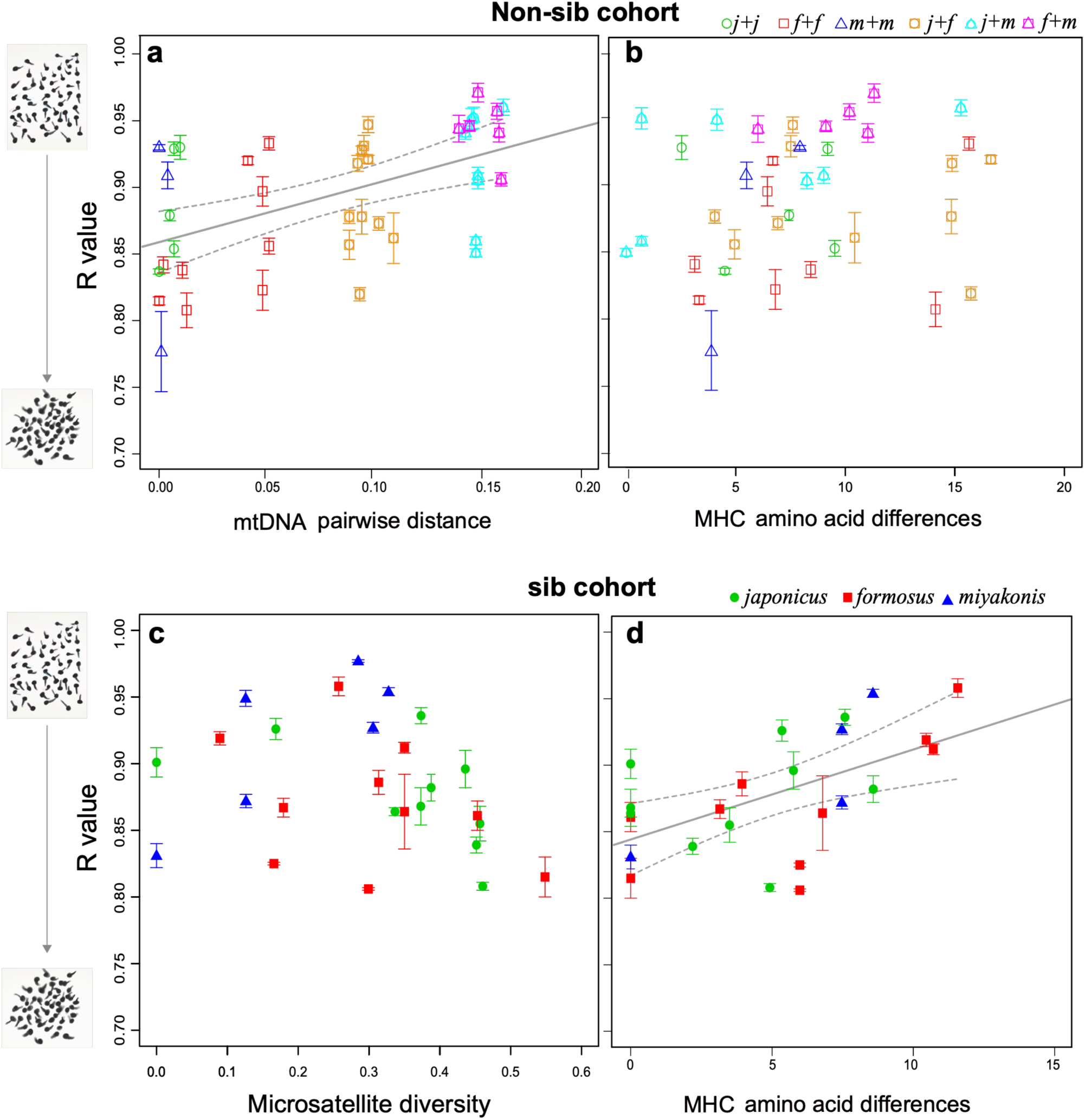
Relationship between aggregation behaviour and genetic diversity at two different scales. The R value represents the degree of aggregation behaviour of each cohort (see text). Regression lines were calculated for significant Spearman’s rank correlation coefficients and the dashed lines indicate two-sided 95% confidence intervals. (**a**) R values and mitochondrial DNA (mtDNA) pairwise distances in non-sib cohorts (*n* = 42, *r* = 0.534; *P* < 0.001; y = 0.3814x + 0.8619). (**b**) R value and major histocompatibility complex (MHC) class II allele amino acid differences in non-sib cohorts (*n* = 38, *r* = 0.231; *P* = 0.163). (**c**) R value and microsatellite diversity in sib cohorts (*n* = 26, *r* = −0.350; *P* = 0.079). (**d**) R value and MHC class II allele amino acid differences in sib cohorts (*n* = 24, *r* = 0.573; *P* = 0.003; y = 0.0058x + 0.8483). Error bars represent standard errors within each cohort for repeated trials. The plot marks indicate the lineage attributes for each cohort tested: green open circles represent non-sib pairs of *japonicus* (= *j* + *j*); red open squares represent non-sib pairs of *formosus* (= *f* + *f*); blue open triangles represent non-sibling pairs of *miyakonis* (= *m* + *m*); overlapping circles and squares in orange represent combinations of *japonicus* and *formosus* lineages (= *j* + *f*); overlapping circles and triangles in aqua represent combinations of *japonicus* and *miyakonis* lineages (= *j* + *m*); overlapping squares and triangles in purple represent combinations of *formosus* + *miyakonis* (= *f* + *m*). Green circles are sib pairs of *japonicus*; red squares are *formosus*; blue triangles are *miyakonis*.

### Scale-dependent associations between genetic variation and aggregation

Model performance evaluations based on the AIC showed a clear difference in the parameters of the candidate models between the sib and non-sib cohorts. For non-sib cohorts, AIC scores determined three candidate models: the best-supported model identified only mtDNA as an explanatory variable for aggregation, whereas the other two models identified combinations of mtDNA and developmental stage or relatedness (Table 2). For sib cohorts (*n* = 26), three candidate models were selected, all of which have MHC as an explanatory variable (Table 2). At both the sib and non-sib level, although there was no model that reached *w_i_* > 0.5, which is often used as a criterion for identifying a good model [51]; the first models were the most parsimonious and possessed the lowest AIC values. Hence, we used these first models on a priority basis; MHC was included in all candidate models for sibs, whereas mtDNA was the only predictor shared by all candidate models for non-sibs (Table 2). In sib cohorts, MHC was correlated with R (t-value = 2.744, *P* = 0.0118). This was supported by the other candidate models (Table 2). In the second the developmental stage and MHC were included as explanatory variable; the developmental stage was not significant (t-value = –0.943, *P* = 0.3562), whereas MHC was significant (t-value = 2.812, *P* = 0.0104). In the third model, microsatellite diversity and MHC were included as explanatory variables; microsatellite diversity was not significant (t-value = –0.781, *P* = 0.4434), whereas MHC was significant (t-value = 2.607, *P* = 0.0165).

**Table 2.** Comparison of the explanatory models for the aggregation index for the sib and non-sib cohorts. The models are ranked according to the Akaike information criterion (AIC): df, degrees of freedom; logLik, log-likelihood; ΔAIC, AIC differences in the model set; wi, Akaike weight. Candidate models (ΔAIC < 2) are shown in bold.

| Model parameters | df | logLik | AIC | ΔAIC | $w_i$ |
| --- | --- | --- | --- | --- | --- |
| **Sib data** |  |  |  |  |  |
| **MHC** | **3** | **44.545** | **-83.1** | **0.00** | **0.210** |
| **Dev + MHC** | **4** | **45.044** | **-82.1** | **1.00** | **0.127** |
| **MHC + Microsat** | **4** | **44.889** | **-81.8** | **1.31** | **0.109** |
| lineage + MHC | 5 | 45.328 | -80.7 | 2.44 | 0.062 |
| Dev + MHC + Microsat | 5 | 45.293 | -80.6 | 2.50 | 0.060 |
| lineage | 4 | 44.226 | -80.5 | 2.64 | 0.056 |
| MHC + Microsat + MHC:Microsat | 5 | 45.165 | -80.3 | 2.76 | 0.053 |
| Null model | 2 | 42.080 | -80.2 | 2.93 | 0.048 |
| Dev + MHC + Microsat + MHC:Microsat | 6 | 45.986 | -80.0 | 3.12 | 0.044 |
| Microsat | 3 | 42.887 | -79.8 | 3.32 | 0.040 |
| lineage + MHC + Microsat | 6 | 45.678 | -79.4 | 3.73 | 0.032 |
| Dev + lineage + MHC | 6 | 45.641 | -79.3 | 3.81 | 0.031 |
| lineage + Microsat | 5 | 44.443 | -78.9 | 4.20 | 0.026 |
| Dev + lineage | 5 | 44.257 | -78.5 | 4.58 | 0.021 |
| lineage | 3 | 42.214 | -78.4 | 4.66 | 0.020 |
| Dev + Microsat | 4 | 42.948 | -77.9 | 5.20 | 0.016 |
| Dev + lineage + MHC + Microsat | 7 | 45.929 | -77.9 | 5.23 | 0.015 |
| lineage + MHC + Microsat + MHC:Microsat | 7 | 45.748 | -77.5 | 5.59 | 0.013 |
| Dev + lineage + Microsat | 6 | 44.457 | -76.9 | 6.18 | 0.010 |
| Full model | 8 | 46.238 | -76.5 | 6.62 | 0.008 |
| **Non-sib data** |  |  |  |  |  |
| **mtDNA** | **3** | **73.900** | **-141.8** | **0.00** | **0.261** |
| **Dev + mtDNA** | **4** | **74.615** | **-141.2** | **0.57** | **0.197** |
| **mtDNA + relatedness** | **4** | **74.200** | **-140.4** | **1.40** | **0.130** |
| Dev + mtDNA + relatedness | 5 | 74.773 | -139.5 | 2.25 | 0.085 |
| mtDNA + lineage | 8 | 77.353 | -138.7 | 3.09 | 0.056 |
| lineage + mtDNA + relatedness | 8 | 77.353 | -138.7 | 3.09 | 0.056 |
| Dev + lineage + mtDNA + relatedness | 9 | 78.254 | -138.5 | 3.29 | 0.050 |
| relatedness | 3 | 71.510 | -137.0 | 4.78 | 0.024 |
| Dev + relatedness | 4 | 72.397 | -136.8 | 5.01 | 0.021 |
| Dev + lineage | 8 | 76.183 | -136.4 | 5.43 | 0.017 |
| Dev + lineage + relatedness | 8 | 76.183 | -136.4 | 5.43 | 0.017 |
| lineage | 7 | 75.159 | -136.3 | 5.48 | 0.017 |
| lineage + relatedness | 7 | 75.159 | -136.3 | 5.48 | 0.017 |
| Null model | 2 | 67.169 | -130.3 | 11.46 | 0.001 |
| Dev | 3 | 67.459 | -128.9 | 12.88 | 0.000 |
| Dev + MHC + mtDNA + MHC | 5 | 67.024 | -124.0 | 17.75 | 0.000 |
| mtDNA + MHC | 4 | 66.012 | -124.0 | 17.78 | 0.000 |
| mtDNA + MHC + relatedness | 5 | 66.935 | -123.9 | 17.93 | 0.000 |
| Dev + mtDNA + MHC + lineage | 10 | 71.718 | -123.4 | 18.36 | 0.000 |
| Dev + mtDNA + MHC + lineage + relatedness | 10 | 71.718 | -123.4 | 18.36 | 0.000 |
| Dev + mtDNA + MHC + relatedness | 6 | 67.633 | -123.3 | 18.53 | 0.000 |
| mtDNA + MHC + lineage + relatedness | 9 | 70.592 | -123.2 | 18.62 | 0.000 |
| mtDNA + MHC + lineage | 9 | 70.592 | -123.2 | 18.62 | 0.000 |
| Dev + mtDNA + MHC + mtDNA:MHC | 6 | 67.475 | -123.0 | 18.85 | 0.000 |
| mtDNA + MHC + mtDNA:MHC | 5 | 66.338 | -122.7 | 19.12 | 0.000 |
| relatedness + mtDNA + MHC + mtDNA:MHC | 6 | 67.330 | -122.7 | 19.14 | 0.000 |
| Dev + relatedness + mtDNA + MHC + mtDNA:MHC | 7 | 68.129 | -122.3 | 19.54 | 0.000 |
| Dev + lineage + MHC | 9 | 69.856 | -121.7 | 20.09 | 0.000 |
| Dev + lineage + MHC + relatedness | 9 | 69.856 | -121.7 | 20.09 | 0.000 |
| Full model | 11 | 71.813 | -121.6 | 20.17 | 0.000 |
| Dev + lineage + mtDNA + MHC + mtDNA:MHC | 11 | 71.813 | -121.6 | 20.17 | 0.000 |
| lineage + mtDNA + MHC + mtDNA:MHC | 10 | 70.650 | -121.3 | 20.50 | 0.000 |
| lineage + mtDNA + MHC + mtDNA:MHC + relatedness | 10 | 70.650 | -121.3 | 20.50 | 0.000 |
| lineage + MHC | 8 | 68.210 | -120.4 | 21.38 | 0.000 |
| lineage + MHC + relatedness | 8 | 68.210 | -120.4 | 21.38 | 0.000 |
| Dev + relatedness + MHC | 5 | 64.655 | -119.3 | 22.49 | 0.000 |
| relatedness + MHC | 4 | 63.486 | -119.0 | 22.83 | 0.000 |
| MHC | 3 | 60.512 | -115.0 | 26.78 | 0.000 |
| Dev + MHC | 4 | 61.114 | -114.2 | 27.57 | 0.000 |

Parameters: mtDNA, mitochondrial DNA pairwise distance; MHC, MHC differences; mtDNA:MHC, interaction effect of mtDNA and MHC; relatedness, sib or intra-/inter-lineage; Dev, developmental stage; lineage, affiliation of mtDNA lineage of tested pair; Microsat, microsatellite genotype diversity; MHC:Microsat, interaction effect of MHC and Microsat.

Meanwhile, in the non-sib cohorts, the GLMs showed that mtDNA was correlated to R value (t-value = 3.888, *P* = 0.0004). This was supported by the other candidate models (Table 2). In the second model, developmental stage and mtDNA were included as explanatory variables; developmental stage was not significant (t-value = 1.162, *P* = 0.2521), whereas mtDNA was significant (t-value = 3.979, *P* = 0.0003). In the third model, relatedness and mtDNA were included as explanatory variables; relatedness was not significant (t-value = 0.750, *P* = 0.4580), whereas mtDNA was significant (t-value = 2.309, *P* = 0.0263).

## DISCUSSION

### Genetic similarity and social aggregation across scales

Our results show that the relationship between genetic variation and tadpole aggregation cannot be reduced to a simple difference between siblings and non-siblings (Figure 2). Although the overall degree of aggregation did not differ significantly among sibling, intra-lineage non-sibling, and inter-lineage non-sibling cohorts, aggregation was related to the genetic variation within both sibling and non-sibling cohorts. Moreover, the genetic variables associated with aggregation differed between these two ranges of relatedness.

Among non-siblings, aggregation decreased with increasing mtDNA pairwise distance, whereas among siblings, aggregation decreased with increasing MHC amino-acid difference (Figure 3).

The absence of a categorical difference between siblings and non-siblings is also consistent with our recent study of *formosus*, in which no kin-biased association was detected in a binary-choice test across different early social and water-rearing conditions [22]. Together, these findings suggest that kin-biased association in *formosus* is not consistently expressed as a categorical preference for siblings over non-siblings.

Importantly, however, the two studies examined different aspects of social behaviour: the previous study tested individual association preference toward sibling versus non-sibling stimuli, whereas the present study quantified the cohesiveness of mixed groups across a broader range of genetic relatedness. The present results therefore extend the absence of a simple kin-biased preference by showing that genetic similarity can nevertheless be related to group cohesiveness at different scales. The additional analysis of spatial distribution provides further support for this interpretation. We found no evidence that cohorts with different relatedness formed distinct spatial subgroups, as assessed by the Morisita index. Thus, the observed relationship between genetic variation and the overall degree of aggregation does not appear to result simply from the formation of separate kin-specific clumps. Instead, genetic composition derived from population genetic structure may be related to the cohesiveness of the social group without necessarily producing discrete assortative groups.

### Different genetic influences at different scales

The different tendencies observed at the two genetic scales raise the possibility that social discrimination is not scale-invariant, but scale-dependent. At the broader scale represented by non-sibling cohorts, mtDNA divergence was associated with aggregation (Figure 3a), whereas at the closer scale represented by sibling cohorts, MHC variation was associated with aggregation (Figure 3d). Importantly, these genetic measures should be regarded as indices of genetic similarity rather than direct evidence of the cues used by tadpoles. mtDNA divergence can covary with variation elsewhere in the genome, and the partial MHC sequence examined here cannot establish whether MHC-derived cues themselves mediate social association. Likewise, the four microsatellite loci used to characterize diversity within sibling cohorts do not provide a genome-wide estimate of relatedness.

Thus, our results identify scale-dependent genetic associations with aggregation, but do not resolve the sensory or molecular mechanisms of social discrimination. We cannot determine how tadpoles assess genetic differences during social interactions, and our genetic indices should therefore be interpreted as correlates of aggregation rather than direct evidence of the cues used by tadpoles.

Greater genetic difference was associated with a lower degree of aggregation, consistent with a positive association between genetic similarity and aggregation behaviour. According to the microsatellite diversity results (Figure 3c), similarities among MHC genes would not necessarily imply similarities among other neutral genes. The relationship between MHC amino-acid differences and aggregation in sibling cohorts is consistent with a possible role of MHC-linked variation in social association, but does not establish MHC-mediated kin recognition. On the other hand, our data did not show a significant association between MHC amino-acid differences and aggregation behaviour in non-sib cohorts (Figure 3b), suggesting that the genetic variables correlated with social aggregation may differ across scales of genetic relatedness.

These findings also suggest that kin discrimination should not necessarily be equated with group favoritism. Although genetic variation was related to the overall cohesiveness of tadpole groups, the Morisita analysis provided no evidence that tadpoles formed distinct kin-specific clumps. Thus, genetic differences may be related to the degree of group cohesion without producing kin-biased association. In this sense, kin discrimination may involve a more continuous relationship with genetic similarity rather than a simple categorical distinction between kin and non-kin.

### Limitations and implications of genetic indices

The mechanisms underlying the observed scale-dependent associations remain unclear. Our genetic indices are insufficient to identify the genetic mechanisms underlying social association. In particular, mtDNA and microsatellite markers provide indirect measures of genetic similarity, while the partial MHC sequence examined here provides only a limited view of MHC variation. Furthermore, the genome-wide relatedness and population genetic structure of the parents contributing to each clutch remain unclear. Genome-wide analyses will therefore be necessary to determine the genetic basis of the observed associations.

Genetic relatedness and kin recognition involve multiple interacting factors [4,18]. In nature, dispersal and life-history processes can constrain the potential for kin-biased social associations [3, 52]. Previous research has also emphasized the context dependence of kin recognition and social preferences [3, 18]. In addition to differences in population genetic structure, breeding sites differ in environmental conditions, including microbial communities and predator regimes, which may contribute to variation in tadpole social behaviour. Consequently, in this study, variation in the partial MHC sequence was associated with the degree of aggregation behaviour in sibling cohorts; however, we could not determine how tadpoles respond to genetic differences among more distantly related individuals.

Our findings therefore suggest that genetic similarity can be associated with social aggregation across multiple scales of relatedness. The relevant genetic scale may matter because genetic similarity among close relatives and more distantly related individuals may be reflected in different genomic and phenotypic variation. Determining whether tadpoles actually use MHC-associated odours, genome-wide similarity, or other correlated traits will require experiments that directly manipulate candidate cues and genomic approaches that quantify relatedness across the genome. Such approaches will be necessary to distinguish genetic correlates of social association from the mechanisms by which tadpoles perceive and respond to one another.

## CONCLUSIONS

Our findings suggest that genetic similarity is related to tadpole group cohesion across multiple scales of genetic relatedness. Determining the cues and mechanisms underlying these relationships will require experiments that directly manipulate candidate cues and assess genetic relatedness at broader genomic scales.

### Ethics

All animals were treated carefully to minimise physical stress. Experiments were performed in accordance with the relevant guidelines of the Life Science Research Ethics and Safety of the University of Tokyo.

## Supporting information

https://drive.google.com/drive/folders/1-v1jKJmNK315HN9XvzNd9V2O3JJH5L4t?usp=drive_link

## Data accessibility

Sequence data are available from GenBank (accession Nos. LC071519–LC071529; LC065649–LC065661). The time-series raw data for all aggregation tests, raw data for analysis of I delta, and haplotype data for all tested tadpoles are provided in the supplementary files.

## Author contributions

K.H prepared the manuscript based on discussions with M.S. K.H. carried out the sampling and the laboratory experiments. M.S.A. analysed the image data.

## Acknowledgements

We thank Prof. Emeritus M. Shimada of the University of Tokyo for suggestions and critical reading of the manuscript. We are grateful for the help of Prof. B. Waldman and Dr. A. Bataille of Seoul National University. We sincerely thank Prof. N. Nikoh of the Open University of Japan and Assoc. Prof. T. Iwasaki of the Ochanomizu University for advice regarding the molecular work. Special thanks are also due to K. Okayama, H. Fujita, K. Higashi, and K. Tanaka, the University Tokyo, Nikko Botanical Garden, and the University of Tokyo, Chichibu Forest for their help with the field work, and Miyata-kun for experimental support.

## Supplementary files are available at the Google Drive

(https://drive.google.com/drive/folders/1-v1jKJmNK315HN9XvzNd9V2O3JJH5L4t?usp=drive_link)

- Experimental Procedure Methods, Supplemental Tables S1,2 and Figures S1–6
- Source data for clutch genotyping
- Source data for MHC haplotype
- Raw data of the aggregation tests

## References

1. Hamilton WD. 1964a The genetical evolution of social behaviour I. J. Theor. Biol. 7, 1–16. (doi:10.1016/0022-5193(64)90038-4)

2. Hamilton WD. 1964b The genetical evolution of social behaviour. II. J. Theor. Biol. 7, 17–52. (doi: 10.1016/0022-5193(64)90039-6)

3. Hepper PG. (Ed.) 2005 Kin recognition. Cambridge University Press.

4. Krupp DB, Lisa MD, Benedict CJ. 2001 Cooperation and Conflict in the Light of Kin Recognition Systems. In The Oxford Handbook of Evolutionary Family Psychology (eds C Salmon C, TK Shackelford), 345–362, Oxford University Press.

5. Waldman B, Adler K. 1979 Toad tadpoles associate preferentially with siblings. Nature 282, 611–613. (doi:10.1038/282611a0)

6. Waldman B. 1982 Sibling association among schooling toad tadpoles: field evidence and implications. Anim. Behav. 30, 700–713. (doi:10.1016/S0003-3472(82)80141-3).

7. O’Hara RK, Blaustein AR. 1982 Kin preference behavior in *Bufo boreas* tadpoles. Behav. Ecol. Sociob. 11: 43–49. (doi: 10.1007/BF00297665)

8. Pfennig DW, Reeve HK, Sherman PW. 1993 Kin recognition and cannibalism in spadefoot toad tadpoles. Anim. Behav. 46, 87–94. (doi: 10.1006/anbe.1993.1164)

9. Hase, K, Kutsukake, Developmental effects on social preferences in frog tadpoles, *Rana ornativentris*. Anim. Behav. 154, 7–16. (doi: 10.1016/j.anbehav.2019.06.001)

10. Hase, K, Kutsukake, N. 2022. Plasticity for the kin and conspecific preferences in the frog tadpoles (*Rana ornativentris*). Anim Cogn 25, 1653–1664. (doi.org/10.1007/s10071-022-01661-1)

11. Waldman B, Rice JE, Honeycutt RL. 1992 Kin recognition and incest avoidance in toads. American Zoologist 32, 18–30. (doi:10.1093/icb/32.1.18)

12. Kenny JS. 1969 Pharyngeal mucous secreting epithelia of anuran larvw. Acta Zool. 50, 143–53.

13. Beiswenger RE. 1975. Structure and function in aggregation tadpoles of American toad, *Bufo Americanus*. Herpetologica 31, 222–233.

14. Watt PJ, Nottingham SF, Young S. 1997 Toad tadpole aggregation behaviour: evidence for a predator avoidance function. Anim. Behav. 54, 865–872. (doi: 10.1006/anbe.1996.0512)

15. Lardner B. 2000 Morphological and life history responses to predators in larvae of seven anurans. Oikos 88, 169–180. (doi: 10.1034/j.1600-0706.2000.880119.x)

16. Wells KD. 2010 The ecology and behavior of Amphibian Larvae. In The ecology and Behavior of Amphibians, 557–598 ,University of Chicago Press. Chap. 12.

17. Hase K, Shimada M. 2014 Female polyandry and size-assortative mating in isolated local populations of the Japanese common toad Bufo japonicus. Biol. J. Linn. Soci. 113 236–242. (doi:10.1111/bij.12339)

18. Breed MD. 2014 Kin and nestmate recognition: the influence of WD Hamilton on 50 years of research. Anim. Behav. 92, 271–279. (doi:10.1016/j.anbehav.2014.02.030)

19. Ostrowski EA, Katoh M, Shaulsky G, Queller DC, Strassmann JE (2008) Kin Discrimination Increases with Genetic Distance in a Social Amoeba. PLoS Biol 6(11). (doi: 10.1371/journal.pbio.0060287)

20. Waldman B. 1988 The ecology of kin recognition. Ann. Rev. Ecol. Syst. 19, 543–571. (doi:10.1146/annurev.es.19.110188.002551)

21. Waldman B. 1991 Kin recognition in amphibians. In Kin Recognition (ed PG Hepper) 162–219, Cambridge Univ. Press.

22. Hase, K. 2026 Social isolation, rather than kinship, influences association behavior in toad tadpoles (*Bufo formosus*). Anim Cogn 29 (doi: 10.1007/s10071-026-02090-0)

23. Matsui M. 1984 Morphometric variation analyses and revision of the Japanese toads (Genus *Bufo*, Bufonidae). Contr. Biol. Lab. Kyoto Univ. 26, 209–428.

24. Kawamura H, Nishioka M, Sumida M, Ryuzaki M. 1990 An electro-phoretic study of genetic differentiation in 40 populations of bufo japonicus distributed in Japan. Sci. Rep. Lab. Amphib. Biol. Hiroshima Univ. 10, 1–51.

25. Nishioka M, Kodama Y, Sumida M, Ryuzaki M. 1990 Genetic relationship among 13 *Bufo* species and subspecies elucidated by the method of electrophoretic analyses. Sci. Rep. Lab. Amphib. Biol. Hiroshima Univ. 10 53–91.

26. Igawa T, Kurabayashi A, Nishioka M, Sumida M. 2006 Molecular phylogenetic relationship of toads distributed in the Far East and Europe inferred from the nucleotide sequences of mitochondrial DNA genes. Mole. Phyl. Evol. 38, 250–260.

27. Manning CJ, Wakeland EK, Potts WK. 1992 Communal nesting patterns in mice implicate MHC gene in kin recognition. Nature 360, 581–584. (doi: 10.1038/360581a0)

28. Penn D, Potts W. 1998 MHC–disassortative mating preferences reversed by cross-fostering. Proc. R. Soc. B 265, 1299–1306. (doi: 10.1098/rspb.1998.0433)

29. Yamazaki K, et al. 1988 Familial imprinting determines H-2 selective mating preferences. Science 240, 1331–1332. (doi: 10.1126/science.3375818)

30. Yamazaki K, et al. 2000 Parent–progeny recognition as a function of MHC odortype identity. Proc. Natl. Acad. Sci. USA 97, 10500–10502. (doi: 10.1073/pnas.180320997)

31. Villinger J, Waldman B. 2008 Self-referent MHC type matching in frog tadpoles. Proc. R. Soc. B 275, 1225–1230. (doi:10.1098/rspb.2008.0022)

32. Villinger J, Waldman B. 2012 Social discrimination by quantitative assessment of immunogenetic similarity. Proc. R. Soc. B 279, 4368–4374. (doi: 10.1098/rspb.2012.1279)

33. Olsson M, Madsen T, Nordby J, Wapstra E, Ujvari B, Wittsell H. 2003 Major histocompatibility complex and mate choice in sand lizards. Proc. R. Soc. B 270, 254–S256. (doi:10.1098/rsbl.2003.0079)

34. Bos DH, Williams RN, Gopurenko D, Bulut Z, Dewoody JA. 2009 Condition-dependent mate choice and a reproductive disadvantage for MHC-divergent male tiger salamanders. Mol. Ecol. 18, 3307–3315. (doi: 10.1111/j.1365-294X.2009.04242.x)

35. Medzhitov R, Janeway CA. 2002 Decoding the patterns of self and nonself by the innate immune system. Science 296, 298–300. (doi: 10.1126/science.1068883)

36. Piertney SB, Oliver MK. 2005 The evolutionary ecology of the major histocompatibility complex. Heredity 96, 7–21. (doi: 10.1038/sj.hdy.6800724)

37. Waldman B. 1984 Kin recognition and sibling association among wood frog (*Rana sylvatica*) tadpoles. Behav. Ecol. Sociobiol. 14, 171–180. (doi: 10.1007/BF00299616)

38. Hase K, Shimada M, Nikoh N. 2012 High degree of mitochondrial haplotype diversity in the Japanese Common Toad *Bufo japonicus* in urban Tokyo. Zool. Sci. 29, 702–708. (doi:10.2108/zsj.29.702)

39. May S, Zeisset I, Beebee TJ. 2011 Larval fitness and immunogenetic diversity in chytrid-infected and uninfected natterjack toad (*Bufo calamita*) populations. Conserv. Genet. 12, 805–811. (doi:10.1007/s10592-011-0187-z)

40. Brede EG, Rowe G, Trojanowski J, Beebee TJC. 2001 Polymerase chain reaction primers for microsatellite loci in the Common Toad *Bufo bufo*. Mol. Ecol. Resour. 1, 308–310. (doi:10.1046/j.1471-8278.2001.00120.x)

41. Hase K, Nikoh N, Shimada M. 2013 Population admixture and high larval viability among urban toads. Ecol. Evol. 3, 1677–1691. (doi:10.1002/ece3.578)

42. Gosner KL. 1960 A simplified table for staging anuran embryos and larvae with notes on identification. Herpetologica 16, 183–190.

43. Clark PJ, Evans FC. 1954 Distance to nearest neighbor as a measure of spatial relationships in populations. Ecology 35, 445–453. (doi:10.2307/1931034)

44. Heupel MR, Simpfendorfer CA. 2005 Quantitative analysis of aggregation behavior in juvenile blacktip sharks. Mar. Biol. 147, 1239–1249. (doi: 10.1007/s00227-005-0004-7)

45. Morisita, M. 1959 Measuring of the dispersion of individuals and analysis of the distribution patterns. Mem. Fac. Sci. Kyushu Univ. Ser. E, 2, 215–235.

46. Tamura K, Nei M, Kumar S. 2004 Prospects for inferring very large phylogenies by using the neighbor-joining method. Proc. Natl. Acad. Sci. USA 101, 11030–11035. (doi:10.1073/pnas.0404206101)

47. Tamura K, et al. 2011 MEGA5: molecular evolutionary genetics analysis using maximum likelihood, evolutionary distance, and maximum parsimony methods. Mol. Biol. Evol. 28, 2731–2739. (doi:10.1093/molbev/msr121)

48. Nei M, Kumar S. 2000 Molecular Evolution and Phylogenetics. Oxford University Press.

49. Excoffier L, Lischer HE. 2010 Arlequin suite ver 3.5: a new series of programs to perform population genetics analyses under Linux and Windows. Mol. Ecol. Resour. 10, 564–567. (doi:10.1111/j.1755-0998.2010.02847.x)

50. R Core Team 2015. R: A language and environment for statistical computing. R Foundation for Statistical Computing, Vienna, Austria. URL https://www.R-project.org/.

51. Burnham KP, Anderson DR. 2003 Model Selection and Multimodel Inference: A Practical Information-Theoretic Approach, 2^nd^ edn. Springer, New York.

52. Jonathan P. Green, Jay M. Biernaskie, Milo C. Mee, Amy E. Leedale. 2024. The Evolution of Kin Discrimination Across the Tree of Life. Annual Review of Ecology, Evolution, and Systematics 55:347–367. (doi.org/10.1146/annurev-ecolsys-102221-051057)

