## Supplementary material for "Scale-dependent associations between genetic similarity and aggregation behaviour in three toad lineages": https://drive.google.com/drive/folders/1-v1jKJmNK315HN9XvzNd9V2O3JJH5L4t?usp=drive_link: Raw data of aggregation test.pdf

### Intra-lineage

### *j+j*

320-1R

trial1 trial2 trial3

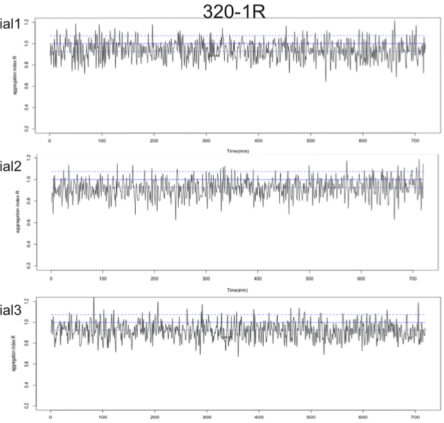

404-1R

trial1 trial2 trial3

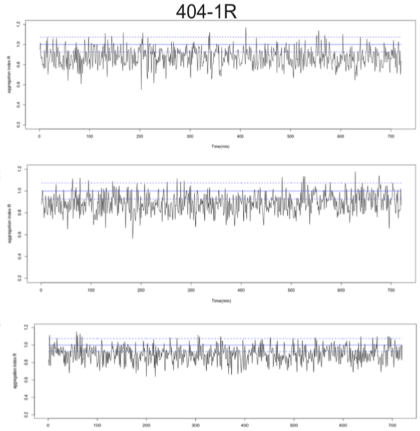

416-1R

trial1 trial2 trial3

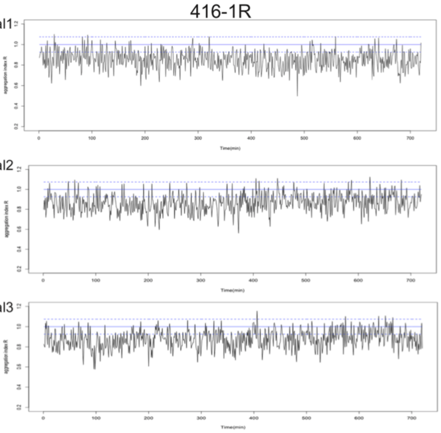

423-2L

trial1 trial2 trial3

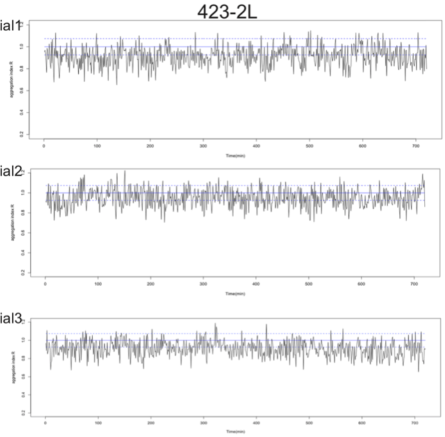

429-2L

trial1 trial2 trial3

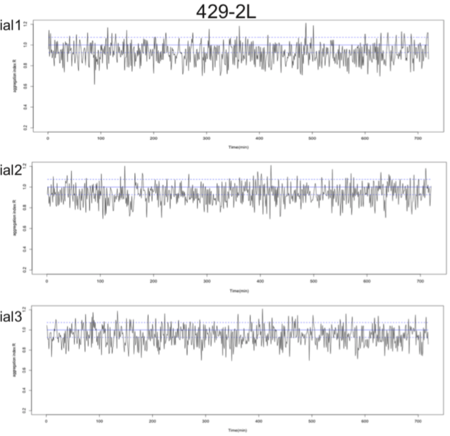

### *f+f*

429-1L

trial1 trial2 trial3

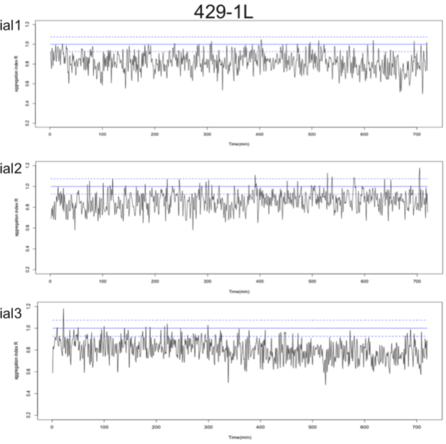

429-2R

trial1 trial2 trial3

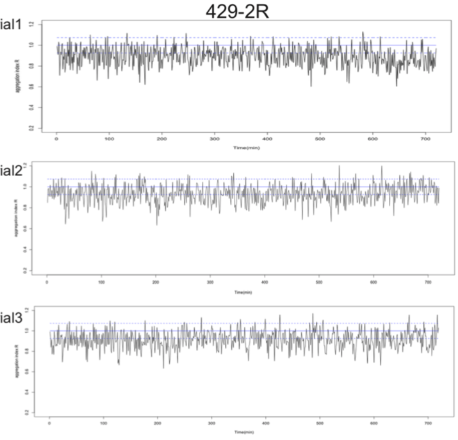

430-1R

trial1 trial2 trial3

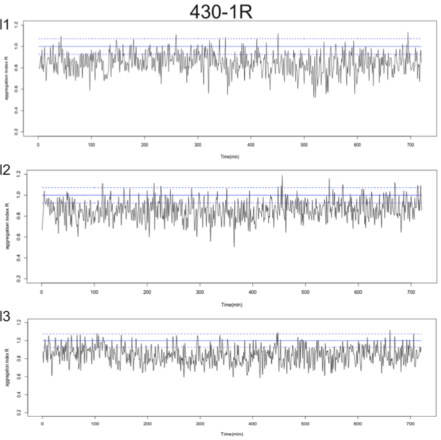

506-2R

trial1 trial2 trial3

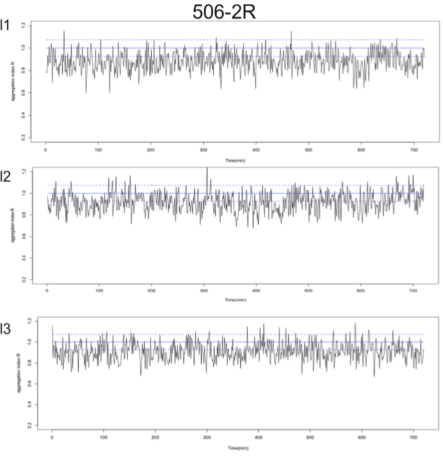

507-1L

trial1 trial2 trial3

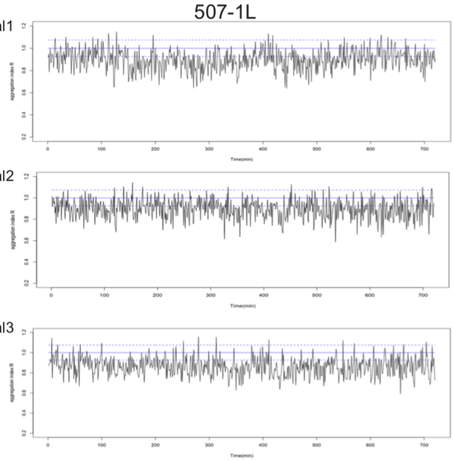

513-1L

trial1 trial2 trial3

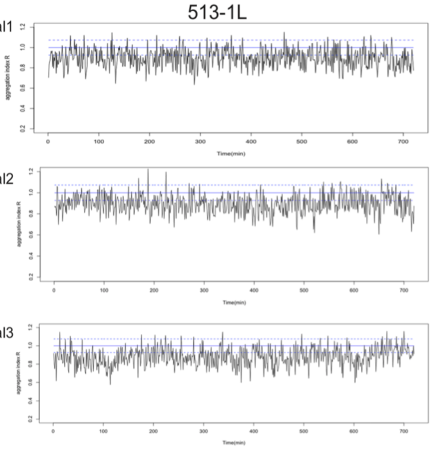

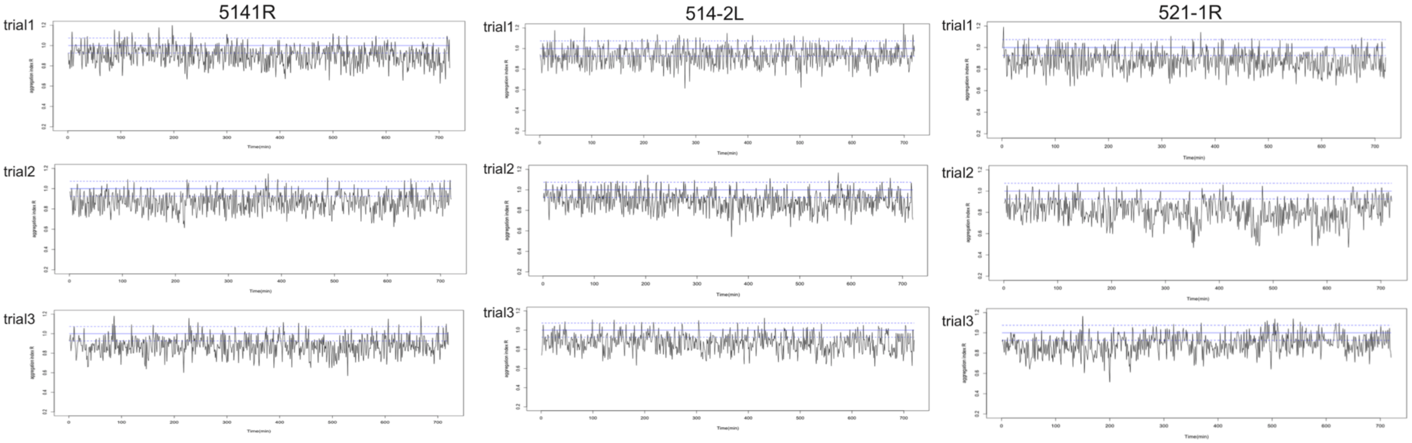


*m+m*

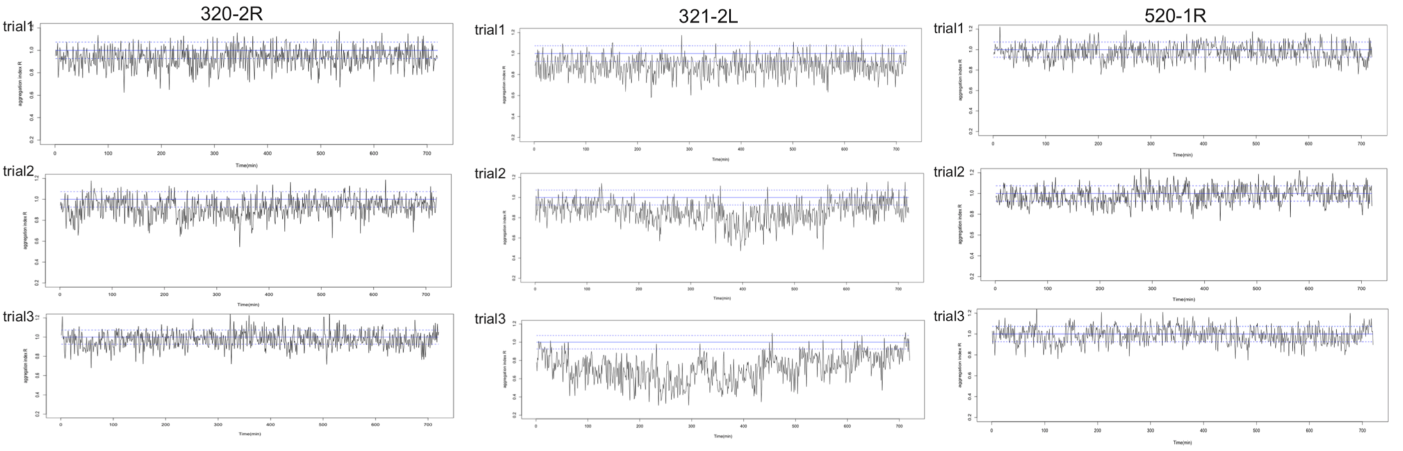


Inter-lineage

*j+f*

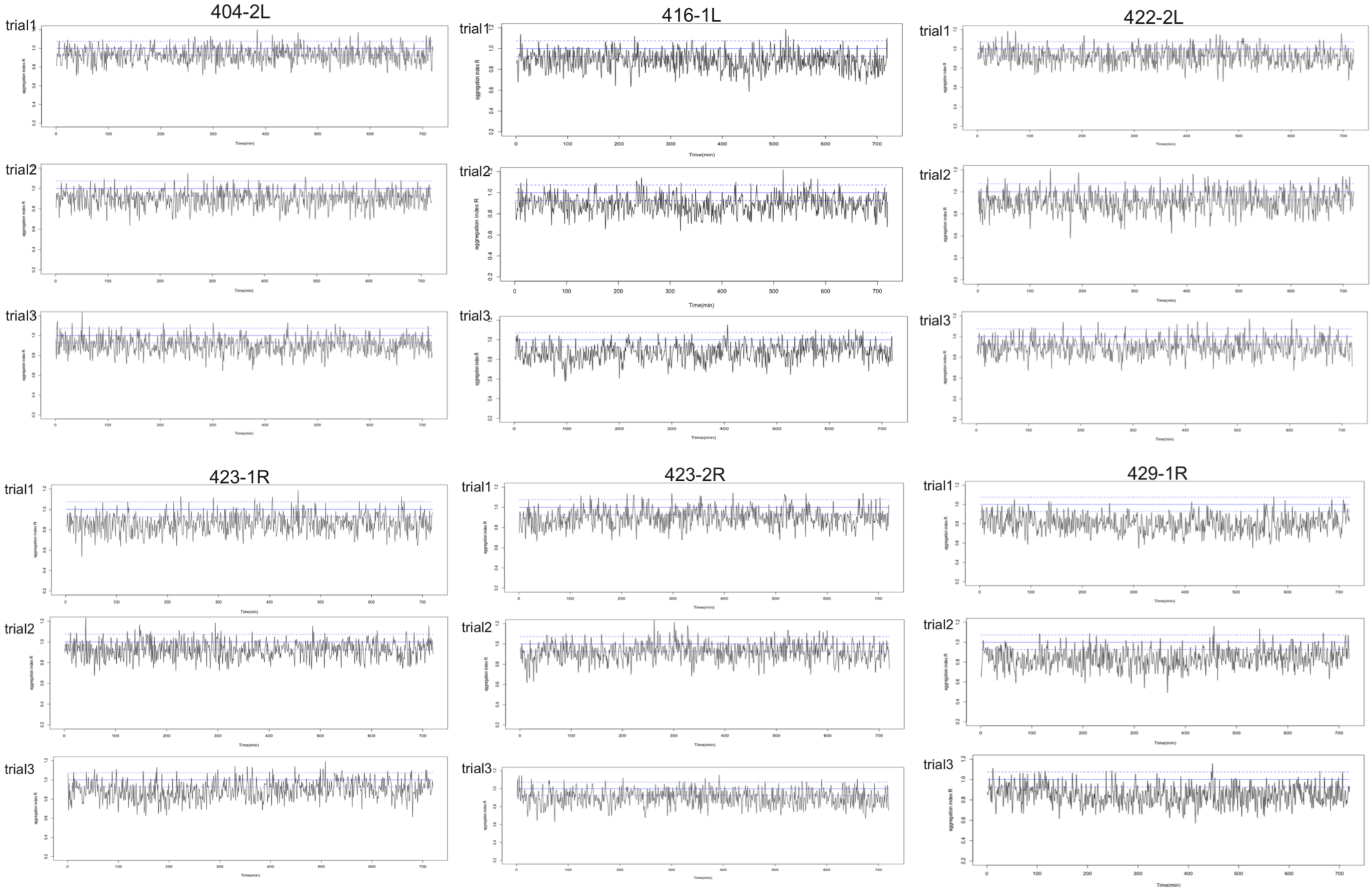

430-2L

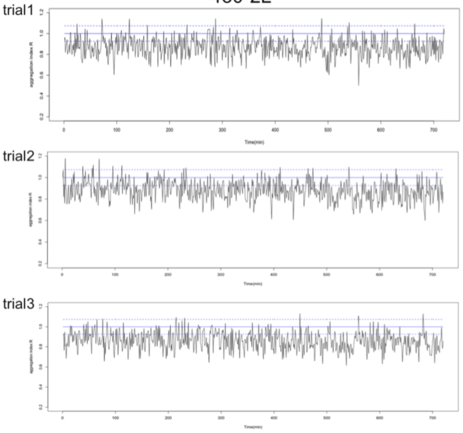


506-1L

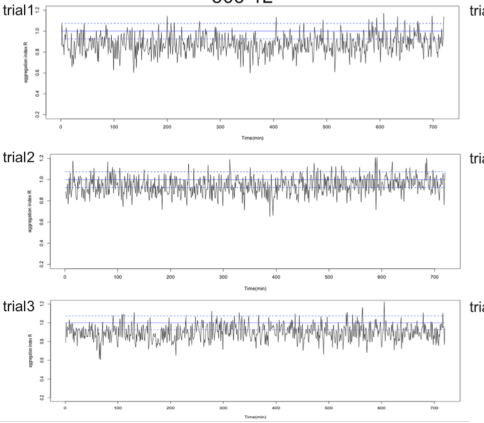


507-2R

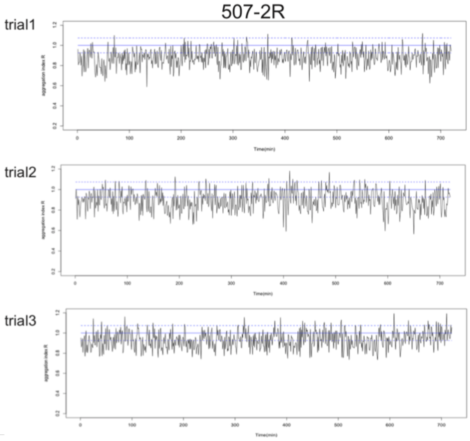


514-2R

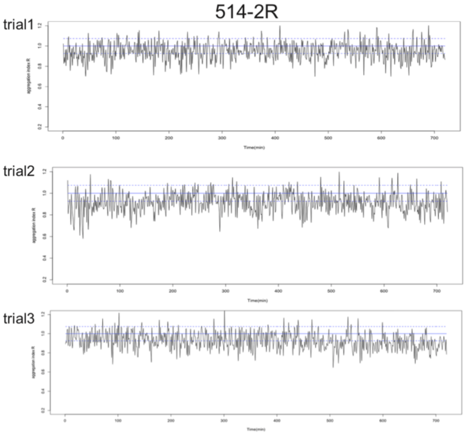


521-1L

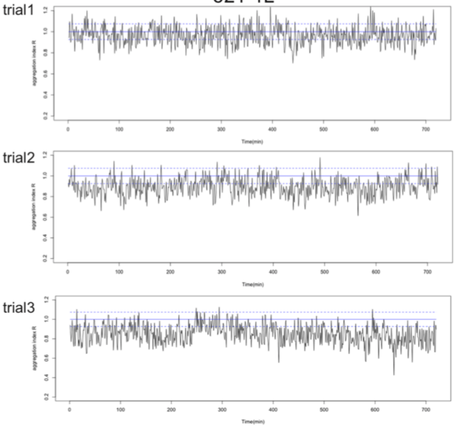


*j+m*

321-1R

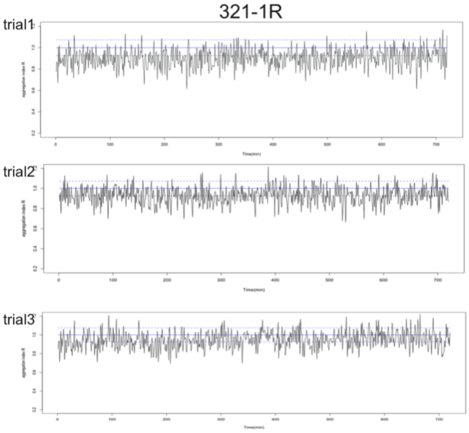


321-2R

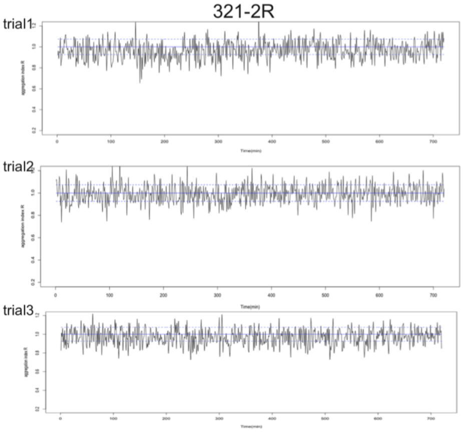


328-1R

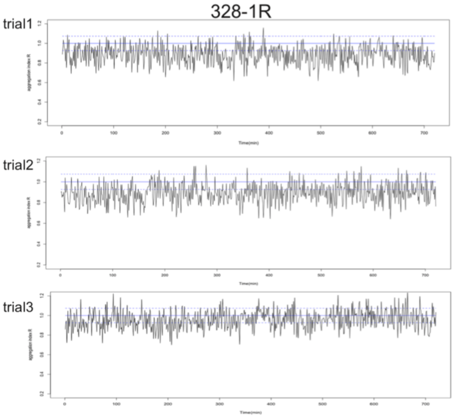


328-2R

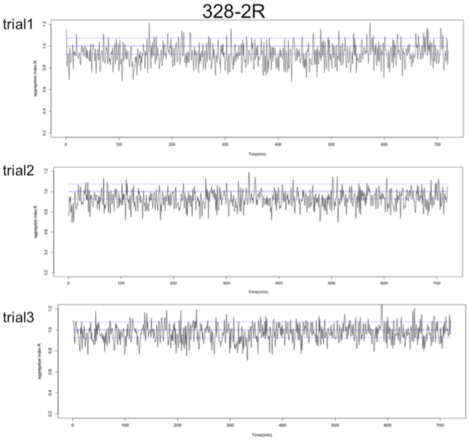


329-2R

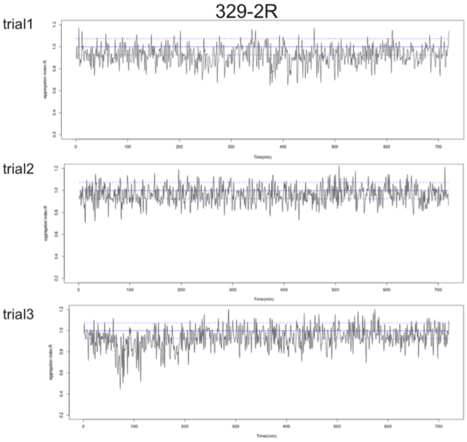


404-1L

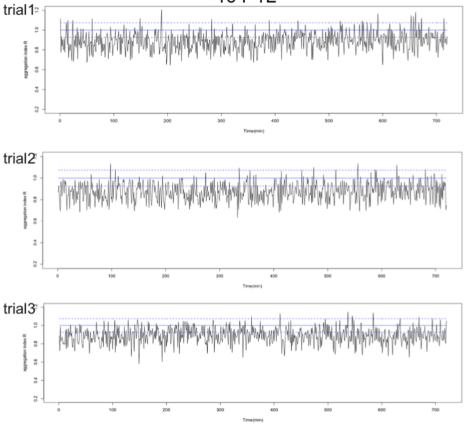

## 404-2R

trial1

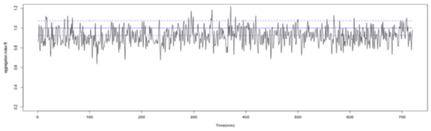

trial2

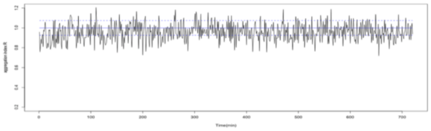

trial3

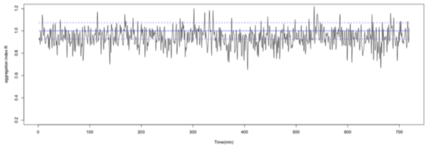

## 520-1L

trial1

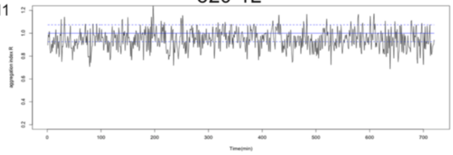

trial2

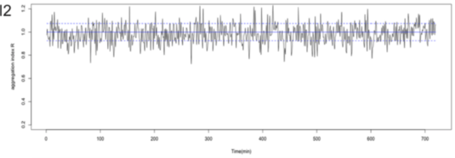

trial3

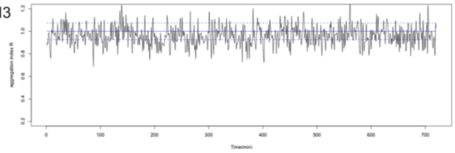

*f+m*

## 422-1L

trial1

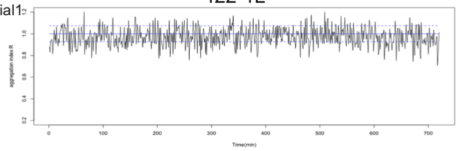

trial2

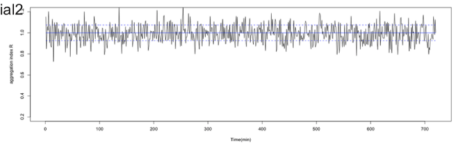

trial3

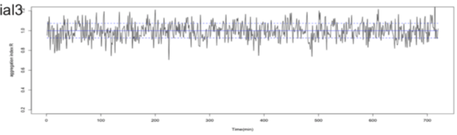

## 422-1R

trial1

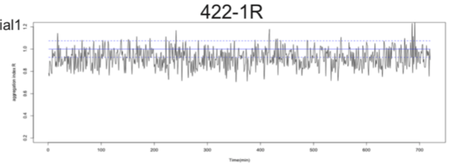

trial2

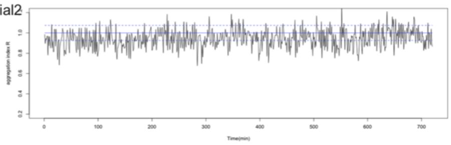

trial3

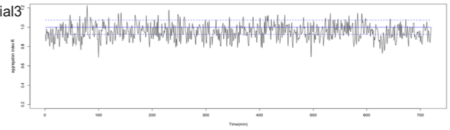

## 430-2R

trial1

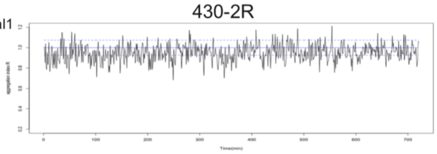

trial2

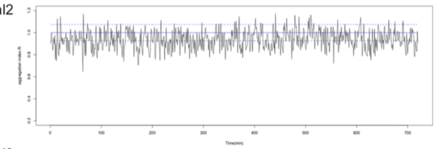

trial3

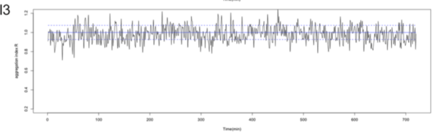

## 507-2L

trial1

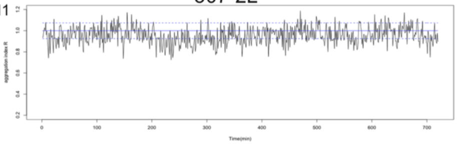

trial2

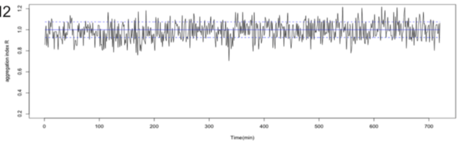

trial3

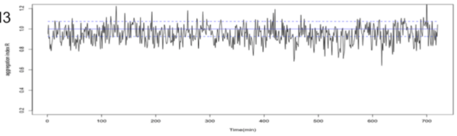

## 513-2L

trial1

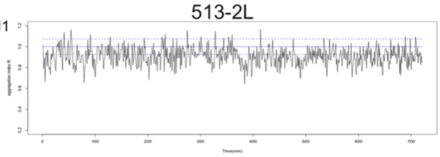

trial2

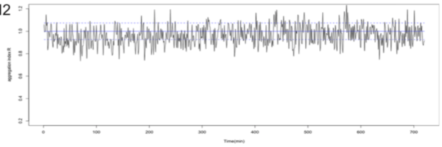

trial3

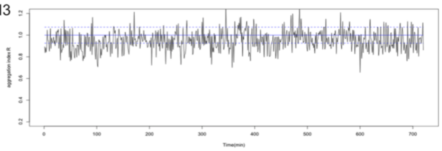

## 513-2R

trial1

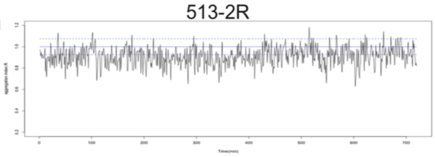

trial2

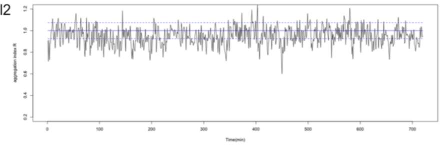

trial3

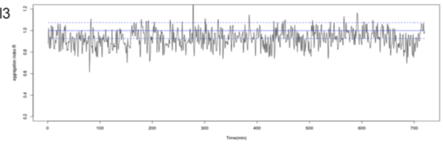
