## Supplementary material for "Scale-dependent associations between genetic similarity and aggregation behaviour in three toad lineages": https://drive.google.com/drive/folders/1-v1jKJmNK315HN9XvzNd9V2O3JJH5L4t?usp=drive_link: Supplementary Method, Tables S1,S2 and Figures.pdf

**Supplementary Method, Tables S1,S2 and Figures S1–S6**

**Experimental Procedures Method (EPM):**

**1. DNA extraction and mtDNA sequencing**

The samples (tadpole's tail tip) were left to digest overnight in ethylenediaminetetraacetic acid-sodium dodecyl sulphate (EDTA-SDS) solution (0.3% SDS, 400 mmol/L NaCl, 5 mmol/L EDTA, 20 mmol/L Tris-HCl, pH 8.0) containing 200 µg/mL proteinase K at 55°C. For sequencing and TA cloning, phenol/chloroform extraction and ethanol precipitation were used. The mtDNA of all clutches was analysed using DNA sequencing, as described previously [S1]; the sequence was used to build a tree in MEGA 5.2 [S2]. Mitochondrial cytochrome b sequences, 0.8 kb in length from 18 clutches, were aligned and deposited in the GenBank nucleotide database [Acc. nos. AB597917, AB597928 from ref. 31; AB713512 from ref. 33; LC071519 – LC071529]. The phylogenetic analyses showed that mtDNA sequences of 18 clutches diverged into three lineages with high bootstrap support (figure S2a).

**2. MHC class II allele**

MHC class II antigen (partial exon) allele sequences were amplified by polymerase chain reaction (PCR) using locus-specific primers developed previously [S3] – 2F347 (GTGACCCTCTGCTCTCCATT) and 2R307b (ATAATTCAGTATATACAGGGTCTCACC) – using KOD FX Neo (Toyobo, Osaka, Japan) in a final reaction volume of 10 µL. The PCR conditions consisted of an initial denaturation step of 2 minutes at 94°C followed by a touchdown program consisting of three cycles of 10 s at 98°C, 30 s at 62°C and 20 s at 74°C, three cycles of 10 s at 98°C, 30 s at 60°C and 20 s at 72°C and 24 cycles of 10 s at 98°C, 30 s at 58°C and 20 s at 70°C. To identify the original alleles within each clutch, 4 – 15 individuals per clutch were sequenced. TA cloning was performed for heterozygous individuals using a pGEM-T vector system (Promega, Madison, WI, USA) and Ligation high (Toyobo) was used for ligation. Transformation was performed using Competent high DH5 (Toyobo). Polyandrous samples were discarded at the sampling stage; therefore, a maximum of four MHC alleles were identified per clutch. Finally, 13 alleles of the 0.3-kb MHC class II locus were detected in 127 individuals from 18 clutches, which were used to construct a phylogenetic tree using MEGA 5.2 [S2]. All MHC alleles have been deposited in the GenBank nucleotide database (Accession Nos. LC065649 – LC065661). From these clutches, 13 MHC alleles presented complex relations with the mtDNA lineages; the BJ1 allele was distributed throughout all lineages (figure S2b). The amino acid alignment of this allele is presented in Figure S3. We then proceeded to genotype all of the tadpoles included in the test (belonging to 68 cohorts). Thus, a total of 3,400 individuals were genotyped using restriction fragment length polymorphism analysis (RFLP) of PCR-amplified fragments (PCR-RFLP).

For individuals belonging to *B. formosus* clutches, we used the reverse primer R_BjMHCII (CCATAGTTG TRTTTACAGWATSTCTCC) during PCR instead of 2R307b, which was developed using already identified alleles. All PCR-amplified MHC sequences were subsequently digested using more than one restriction enzyme, such as *Bme*T1101, *Bse*RI, *Bsg*I or *Xho*II, chosen to distinguish haplotypes among all the possible combinations of alleles identified in the clutches (tables S1 and S2). The MHC haplotypes of all 50 individuals in each cohort could be identified from PCR-RFLP data. However, in cases where the haplotype of more than two individuals could not be identified using this method, the entire cohort was excluded from the genetic diversity analysis. A total of 3,000 individuals (62 of 68 cohorts) were finally included in the analysis.

**3. Microsatellite marker**

To estimate genetic diversity in the sib group, we selected four microsatellite loci by PCR: 1) *Bbufu23*; 2) *Bbufu39*; 3) *Bbufu62* [S4]; and 4) *Bjap14* [S5]. Following the method described previously [S5], the loci were amplified with two multiplex reactions using a Multiplex PCR Kit (QIAGEN, Valencia, CA, USA) according to the manufacturer’s protocol. PCR amplification consisted of an initial denaturation step at 95°C for 15 minutes, followed by 25–35 cycles of 30 s at 94°C for denaturation, primer-specific touchdown annealing for 90 s at 46–61°C [S5] and an extension step at 72℃ for 1 minute. The fragment sizes of the PCR products from each microsatellite locus were analysed using a CEQ8000 Genetic Analysis System (Beckman Coulter, Fullerton, CA, USA) with a Genomelab Size Standard Kit 400 (60 – 420 bp; Beckman Coulter). Overall, we genotyped 1,200 individuals belonging to 42 sib cohorts. A polyandry check was performed in 18 clutches using GERUD2.0 [S6], which is a useful program when information is available regarding the parental genotypes. The results indicated that 17 of 18 clutches had a single sire. One clutch, NK1, was sired by at least two males; the relative paternity results for NK1 showed that the first male sired 84% of the clutch, and the second male sired the remainder. Hence, the word ‘sibs’ does not refer to full-sibs in this paper.

### Supplemental tables and figures:

**Table S1**

Information for the 18 clutches used in the experimental trials. Mitochondrial lineages, sampling site, locality, coordinates, identified MHC class II allele type, number of sequencing samples (figures in parentheses indicate TA cloning samples) used to detect the alleles and RFLP enzymes used for genotyping. To genotype one cohort, we separated 50 tadpoles into two sets (1 – 25 and 26 – 50) and used combinations of one to four of the enzymes listed in the table, considering the clutches from which the tested tadpoles were derived. Thirteen alleles, BJ1 – 13, were detected from among the 18 clutches. Allele BJ1 was present across all lineages, whereas BJ6 – 10 and BJ 3 – 5, 11 and 12 were derived only from *japonicus* and *formosus*, respectively. For the information of site geography, please refer to Figure S1.

| lineage | site | clutch | Locality | Coordinates | MHC allele | | | sequencing | RFLP enzyme |
|---|---|---|---|---|---|---|---|---|---|
| *miyakonis* | **1*** | **DT1** | Minami-Daitō Is., Okinawa | N25.844°E131.229° | BJ1 | BJ2 | | 5(5) | BseRI, BsgI |
| | | **DT2** | Minami-Daitō Is., Okinawa | N25.844°E131.229° | BJ1 | BJ2 | | 6(4) | BseRI, BsgI |
| | | **DT4** | Minami-Daitō Is., Okinawa | N25.835°E131.236° | BJ1 | | | 7(6) | BseRI |
| *japonicus* | **2** | **YW1** | Yoshiwa, Hiroshima | N34.457°E132.132° | BJ1 | BJ6 | | 4(4) | BseRI, BsgI |
| | | **YW2** | Yoshiwa, Hiroshima | N34.457°E132.132° | BJ1 | | | 7(7) | BseRI |
| | | **YW3** | Yoshiwa, Hiroshima | N34.457°E132.132° | BJ1 | BJ7 | BJ8 | 5(3) | BseRI, BmeT1101, XhoII |
| | | **YW4** | Yoshiwa, Hiroshima | N34.457°E132.132° | BJ1 | BJ9 | | 4(3) | BseRI, XhoII |
| | **3** | **OM** | Omogo, Ehime | N33.715°E133.069° | BJ1 | BJ10 | | 4(3) | BseRI, BsgI |
| | **4*** | **KM1** | Komaba, Tokyo | N35.659°E139.688° | BJ1 | | | 12(10) | BseRI |
| | | **KM2** | Komaba, Tokyo | N35.659°E139.688° | BJ1 | BJ2 | | 4(4) | BseRI, BsgI |
| | | **KM3** | Komaba, Tokyo | N35.659°E139.688° | BJ1 | | | 10(9) | BseRI |
| *formosus* | **5** | **NK1** | Nikko, Tochigi | N36.751°E139.586° | BJ1 | BJ3 | BJ13 | 15(10) | BseRI, BmeT1101, BsgI |
| | | **NK2** | Nikko, Tochigi | N36.751°E139.586° | BJ1 | BJ3 | | 10(6) | BseRI, BmeT1101 |
| | **6** | **CB** | Yorii, Saitama | N36.116°E139.220° | BJ3 | BJ11 | BJ12 | 4(3) | BmeT1101, BsgI |
| | **7** | **ES1** | Chichibu, Saitama | N35.888°E138.811° | BJ1 | BJ5 | | 7(7) | BseRI, BmeT1101 |
| | | **ES2** | Chichibu, Saitama | N35.888°E138.811° | BJ1 | BJ5 | | 6(3) | BseRI, BmeT1101 |
| | **8** | **RS1** | Hiki, Saitama | N36.037° E132.294° | BJ4 | | | 12(7) | BsgI |
| | | **RS2** | Hiki, Saitama | N36.037° E132.294° | BJ1 | BJ4 | | 5(3) | BseRI, BsgI |

*Artificially introduced

**Table S2**

Restriction enzymes used for PCR-RFLP to genotype 13 alleles of the MHC class II gene. Numbers indicate restriction fragment sizes, e.g. *Bse*R1 cuts the BJ1 allele into fragments of 0.12 kb and 0.20 kb.

| **Enzyme** | **Recognition site** | **identified MHC alleles used for genotyping** | | | | | | | | | | | | | |
|---|---|---|---|---|---|---|---|---|---|---|---|---|---|---|---|
| | | BJ1 | BJ2 | BJ3 | BJ4 | BJ5 | BJ5 | BJ6 | BJ7 | BJ8 | BJ9 | BJ10 | BJ11 | BJ12 | BJ13 |
| BmeT1101 | c/ycgrg | | | 169/151 | | 169/151 | 169/151 | | 169/151 | | 169/151 | 169/151 | | 169/151 | |
| BseR1 | gaggag | 120/200 | | 120/200 | 120/200 | | | 120/200 | 120/200 | 120/200 | 120/200 | 120/200 | 120/200 | 120/200 | 120/200 |
| BsgI | gtgcag | | 96/224 | | 187/133 | 96/224 | 96/224 | 187/133 | | 187/133 | 96/224 | 96/224 | 96/224 | 96/224 | 187/133 |
| XhoII | r/gatcy | | | | | | | | | 207/113 | 207/113 | | | | |

**Figure S1**

Map of sampling sites. Each number of sites is corresponding to Table S1. Shaded portion indicates natural distribution area based on the previous study [S7].

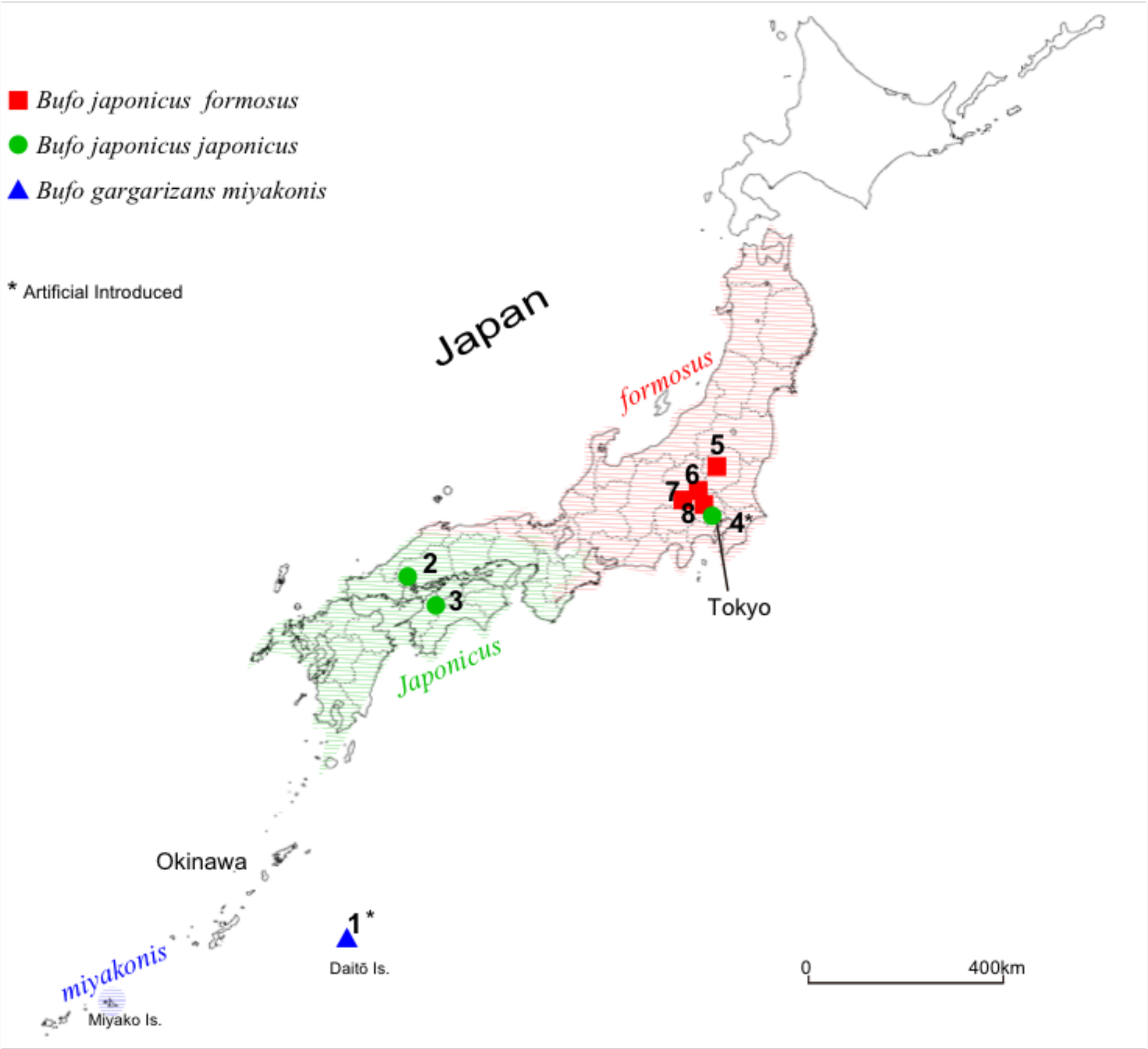

**Figure S2**

Examples of the analysis of $I_\delta$ [S8]. Each image showing the highest value of R has been divided into 16 sections. We calculated $I_\delta$ based on the number of tadpoles in each section. When a tadpole was located in two or more sections, we chose the section containing the greatest part of the tadpole's body.

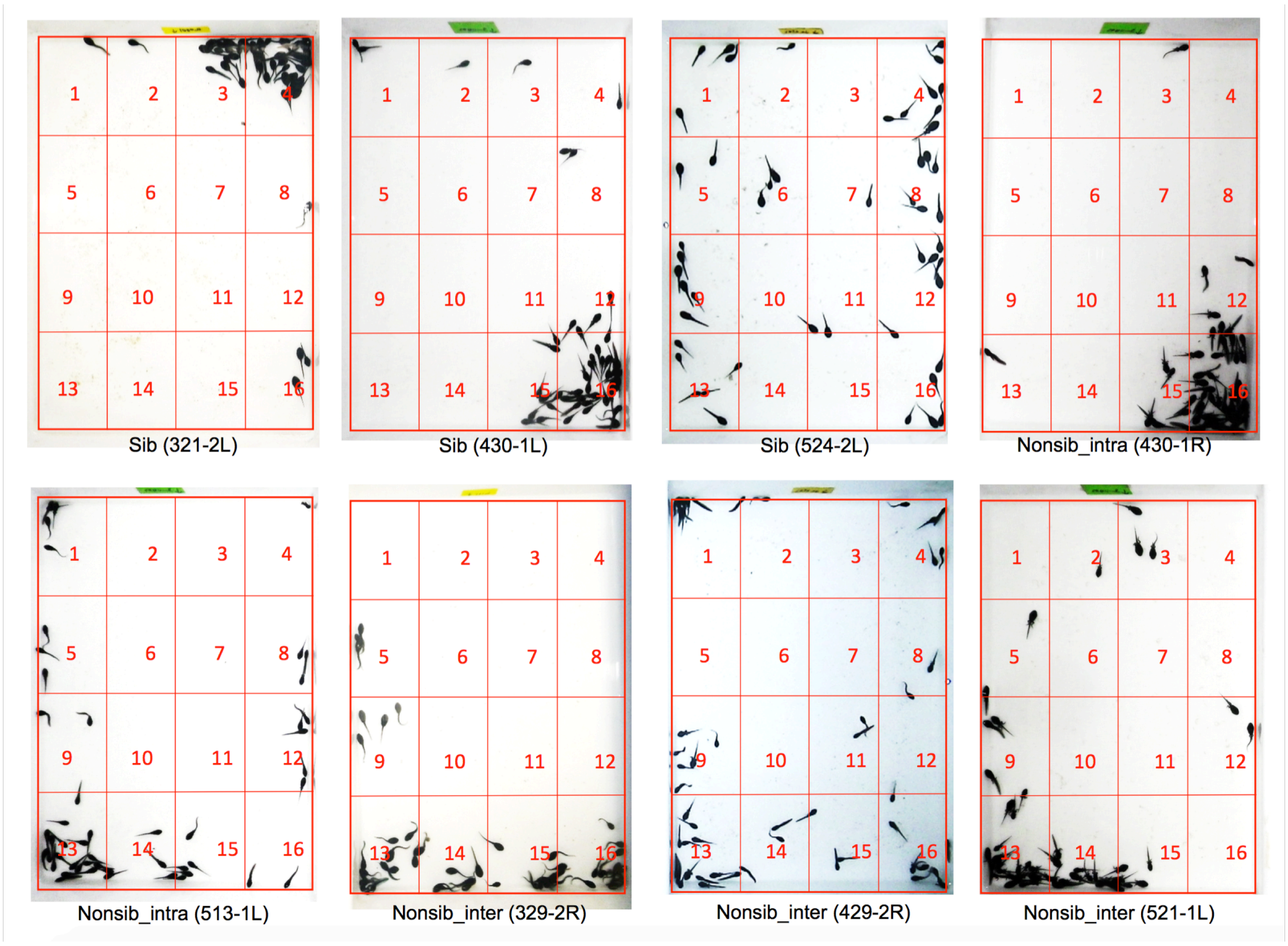


**Figure S3**

Maximum likelihood (ML) trees of the mitochondrial DNA lineage of 18 clutches based on 831 bp of Cytb sequences (**a**) and MHC alleles based on 324 bp of class ll exon partial sequences (**b**). The width of the colour bar on the right side of each allele ID corresponds to the number of identified clutches. The colour represents attribution of the mtDNA lineage of the identified clutches corresponding to the mtDNA tree: green, red and blue indicate, *japonicus*, *formosus* and *miyakonis* lineages, respectively. From 18 clutches, 13 MHC alleles presented complex relations with the mtDNA lineages; the BJ1 allele was distributed throughout all lineages. Both trees were estimated using the GTR +Γ model, selected according to the Akaike information criterion (AIC) value in MEGA 5.0 [S2]. Support values for the internal nodes were inferred from 1,000 bootstrap replicates. †1 and †2 correspond to haplotypes from [S1] and [S5], respectively.

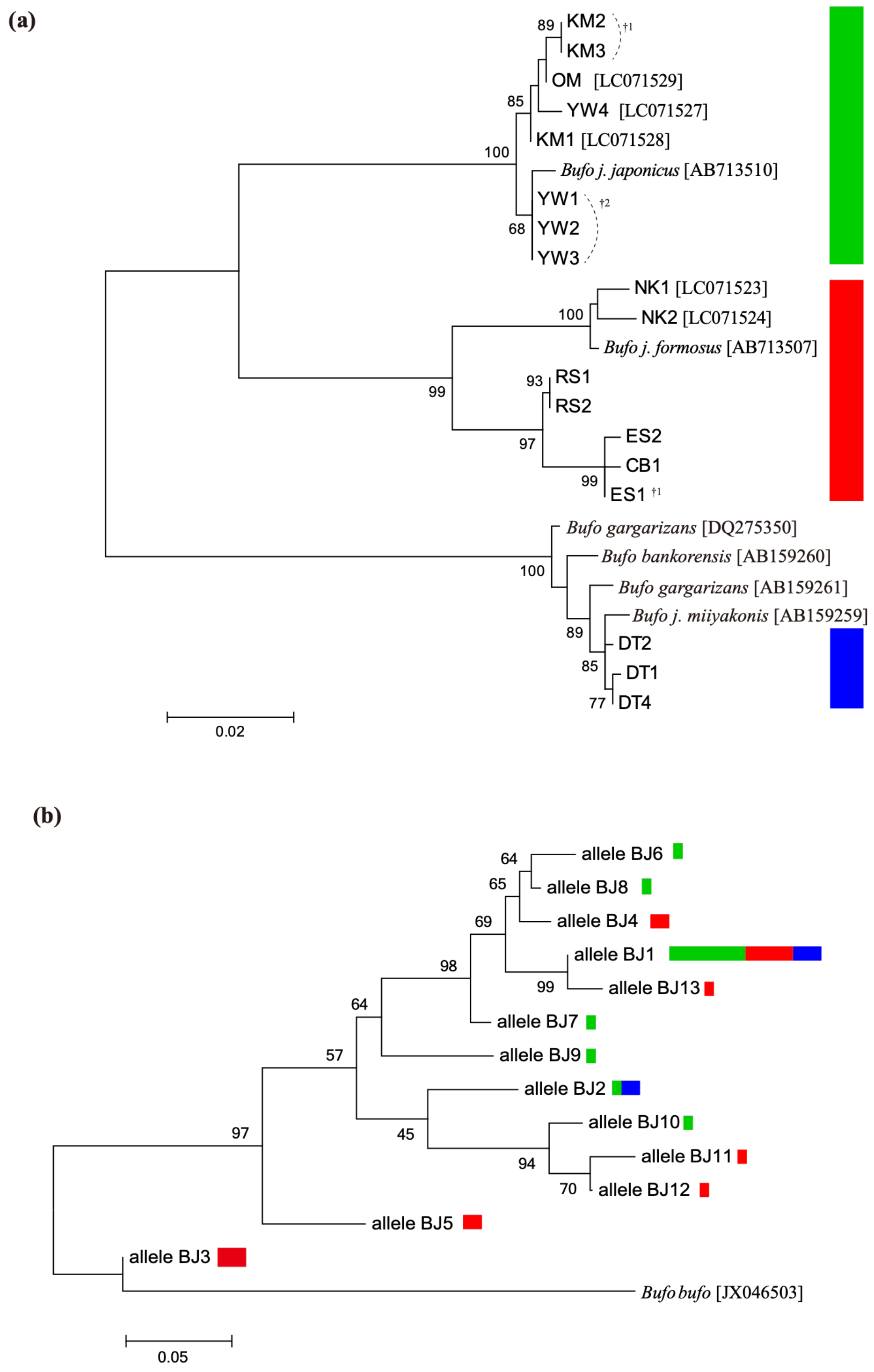

**Figure S4**

Plot of the relationship between the R value and $I_\delta$ ($n$ = 68). A low value of R indicates a high degree of aggregation, whereas a high value of $I_\delta$ indicates low dispersion [S8].

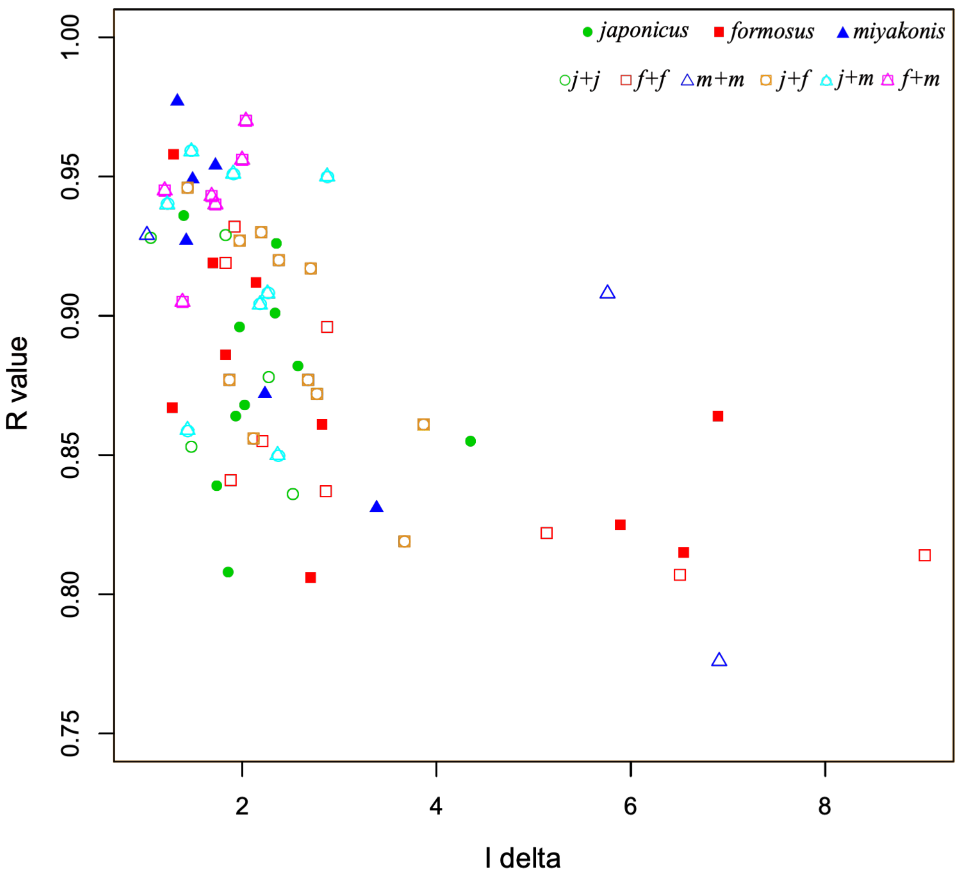


**Figure S5**

Plots of one-way relationships between aggregation score and genetic diversity. Overall, aggregation behavior was significantly correlated with both mtDNA and MHC amino acid differences. Regression lines were calculated for significant Spearman's rank correlation coefficients and the dashed lines indicate two-side 95% confidence intervals. (**a**) R index and mtDNA pairwise distances ($n$ = 68) ($r$ = 0.311, $P$ = 0.010; y = 0.293x + 0.875). (**b**) Aggregation time with lower confidence limits (5%) and mtDNA pairwise distances ($n$ = 68) ($r$ = –0.302, $P$ = 0.012; y = –824.78 + 335.44x). (**c**) R index and MHC amino acid differences ($n$ = 62) ($r$ = 0.371, $P$ = 0.003; y = 0.0012x + 0.869). (**d**) Aggregation time with lower confidence limits (5%) and MHC ($n$ = 62) ($r$ = -0.376, $P$ = 0.002; y = –3.590x + 353.067). Error bars represent standard errors within each cohort for repeated trials. The plot marks indicate the lineage attributes for each cohort tested: green filled circles represent sibling pairs of *japonicus* (= *j* + *j*_sib); red filled squares represent sibling pairs of *formosus* (= *f* + *f*_sib); blue filled triangles represent *miyakonis* (= *m* + *m*_sib); green open circles represent non-sibling pairs of *japonicus* (= *j* + *j*); red open squares represent non-sibling pairs of *formosus* (= *f* + *f*); blue open triangles represent non-sibling pairs of *miyakonis* (= *m* + *m*); overlapping circles and squares in orange represent combinations of *japonicus* and *formosus* lineages (= *j* + *f*); overlapping circles and triangles in aqua represent combinations of *japonicus* and *miyakonis* lineages (= *j* + *m*); overlapping squares and triangles in purple represent combinations of *formosus* + *miyakonis* (= *f* + *m*).

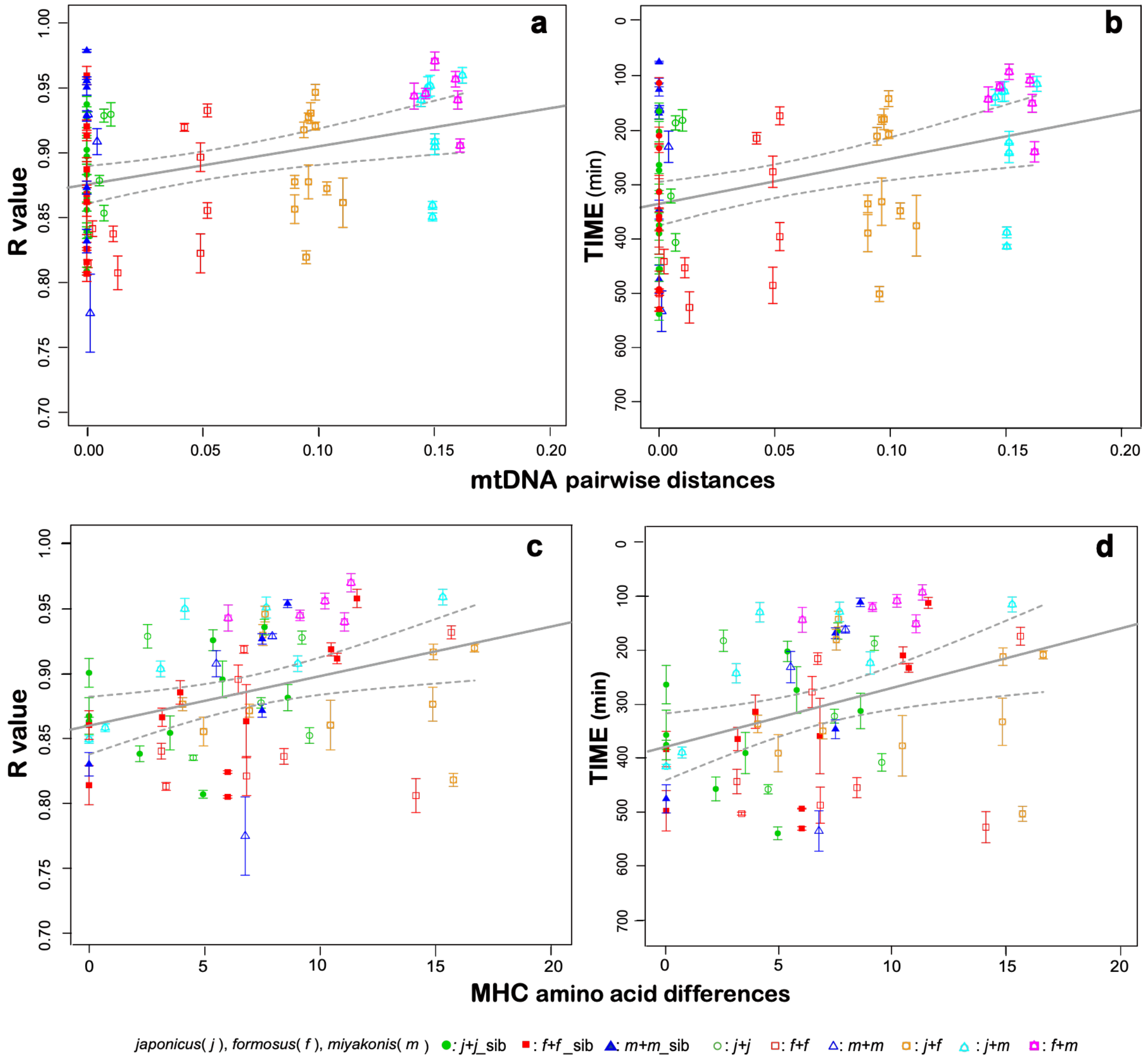

**Figure S6**

Amino acid alignment of the MHC class II partial exon (324 bp) of the 13 identified alleles.

```
allele_BJ1    PLLSITDAVDYMIEVKSECHYVNGTQQVRYLQRYSYNQEEFVYFDSEEGR    50
allele_BJ2    PLLSITDAVDYMTESKFECHYVNGTQQVRFLHRCFYNQEEIVYFDSEEGR    50
allele_BJ3    PLLSITAAVDYMTEVKSECHYVNGTQQVRYLDRYFYNQEELVYFDSEEGR    50
allele_BJ4    PLLSITDAVDYMTEIKSDCHYVNGTQQVRYLQRYSYNQEEFVYFDSEEGR    50
allele_BJ5    PLLSITDAVDYMTEIKNECHYVNGTQQVRYLHRYFYNQEETLYFDSDKGR    50
allele_BJ6    PLLSITDAVDYMIEVKSDCHYVNGTQQVRYLQRYSYNQEEFVYFDSEEGR    50
allele_BJ7    PLLSITDAVDYMIEMKSDCHYVNGTQQVRYLQRYSYNQEEIVYFDSEEGR    50
allele_BJ8    PLLSITDAVDYMIEVKSDCHYVNGTQQVRYLQRYSYNQEEFVYFDSEEGR    50
allele_BJ9    PLLSITDAVDYMTEIKSDCHFLNGTQQVRFLHRYFYNQEELVYFDSEEGR    50
allele_BJ10   PLLSITAAVDYMTEEKFECHYVNGTQQVRYLHRCFYNQEELVYFDSDKGY    50
allele_BJ11   PLLSITAAVDYMTEEKFECHYVNGTQQVRYLHRCFYNQEEIVYFDSDKGR    50
allele_BJ12   PLLSITAAVDYMTEEKFECHYVNGTQQVRYLHRCFYNQEEFVYFDSDKGR    50
allele_BJ13   PLLSITDAVDYMIEVKSECHYVNGTQQVRYLQRYSYNQEEFVYFDSEEGR    50

allele_BJ1    FIAKTEFGKPSADYWNKNKDFIEDRKSAVERFCKHNYG-VFHSVTADRKG    99
allele_BJ2    YIAKTEFGKPDADYWNKNKEIIEQRKSYVETYCKHNYG-VFHSVTADRRG    99
allele_BJ3    YIAKTELGKPDAEAWNKNKEIIEDRKSSVETFCKYNYGTATTVGITGRRG   100
allele_BJ4    FIAKTEFGKPSADYWNKDKDFIEQKKSAVETFCKHNYG-VFHSVTADRRG    99
allele_BJ5    YIAKTELGKPEAEAWNKDKEIIEDRKSAVE?FCKYNYG-VIHSFTADRRG    99
allele_BJ6    FIAKTEFGKPSAELWNKDKDFIEQKKSAVETFCKHNYG-VLHSVTADRRG    99
allele_BJ7    FIAKTELGKPDADYWNKNKDFIEQRKSAVETFCKHNYG-VFHSVTADRRG    99
allele_BJ8    FIAKTEFGKPSADYWNKDKDLIEQRKSAVETFCKHNYG-VLHSVTADRRG    99
allele_BJ9    YIAKTELGKPDADYWNKDKDLIEREKSYVETFCKYNYG-VFHSVTADRRG    99
allele_BJ10   YIAKTELGKPDADYWNKDKEIIEHKKSEVETFCKHNYG-VLHGVTGDRRG    99
allele_BJ11   YIAKTEFGKPDAEAWNKDKEFIEQKKSAVERFCKHNYG-VLHGVTGDRRG    99
allele_BJ12   YIAKTELGKPDAEAWNKDKEIIEQKKSEVETFCKHNYG-VLHGVTGDRRG    99
allele_BJ13   IIAKTEFGKPSADYWNKNKDFIEDRKSAVERFCRHNYG-LYHSVTADRKG    99

allele_BJ1    ETLYILNY                                             107
allele_BJ2    ETLYILNY                                             107
allele_BJ3    ETLYILNY                                             108
allele_BJ4    ETLYILNY                                             107
allele_BJ5    ETLYILNY                                             107
allele_BJ6    ETLYILNY                                             107
allele_BJ7    ETLYILYY                                             107
allele_BJ8    ETLYILNY                                             107
allele_BJ9    ETLYILNY                                             107
allele_BJ10   EPLYILYY                                             107
allele_BJ11   ETLYILNY                                             107
allele_BJ12   ETLYILNY                                             107
allele_BJ13   ETLYILNY                                             107
```

**References**

S1. Hase K, Shimada M, Nikoh N. 2012 High degree of mitochondrial haplotype diversity in the Japanese Common Toad *Bufo japonicus* in urban Tokyo. *Zool. Sci.* **29**, 702–708. (doi:10.2108/zsj.29.702)

S2. Tamura K, et al. 2011 MEGA5: molecular evolutionary genetics analysis using maximum likelihood, evolutionary distance, and maximum parsimony methods. *Mol. Biol. Evol.* **28**, 2731–2739. (doi:10.1093/molbev/msr121)

S3. May S, Zeisset I, Beebee TJ. 2011 Larval fitness and immunogenetic diversity in chytrid-infected and uninfected natterjack toad (*Bufo calamita*) populations. *Conserv. Genet.* **12**, 805–811. (doi:10.1007/s10592-011-0187-z)

S4. Brede EG, Rowe G, Trojanowski J, Beebee TJC. 2001 Polymerase chain reaction primers for microsatellite loci in the Common Toad *Bufo bufo*. *Mol. Ecol. Resour.* **1**, 308–310. (doi:10.1046/j.1471-8278.2001.00120.x)

S5. Hase K, Nikoh N, Shimada M. 2013 Population admixture and high larval viability among urban toads. *Ecol. Evol.* **3**, 1677–1691. (doi:10.1002/ece3.578)

S6. Jones AG. 2005 GERUD 2.0: a computer program for the reconstruction of parental genotypes from half-sib progeny arrays with known or unknown parents. *Mol. Ecol. Notes* **5**, 708–711. (doi:10.1111/j.1471-8286.2005.01029.x)

S7. Matsui M. 1984 Morphometric variation analyses and revision of the Japanese toads (Genus *Bufo*, Bufonidae). *Contr. Biol. Lab. Kyoto Univ.* **26**, 209–428.

S8. Morisita, M. 1959 Measuring of the dispersion of individuals and analysis of the distribution patterns. *Mem. Fac. Sci. Kyushu Univ. Ser.* E, **2**, 215–235.
